# Argonaute Binding within Human Nuclear RNA and its Impact on Alternative Splicing

**DOI:** 10.1101/2021.02.09.430489

**Authors:** Yongjun Chu, Shinnichi Yokota, Jing Liu, Audrius Kilikevicius, Krystal C. Johnson, David R. Corey

## Abstract

Mammalian RNA interference (RNAi) is often linked to the regulation of gene expression in the cytoplasm. Synthetic RNAs, however, can also act through the RNAi pathway to regulate transcription and splicing. While nuclear regulation by synthetic RNAs can be robust, a critical unanswered question is whether endogenous functions for nuclear RNAi exist in mammalian cells. Using enhanced crosslinking immunoprecipitation (eCLIP) in combination with RNA sequencing (RNAseq) and multiple *AGO* knockout cell lines, we mapped AGO2 protein binding sites within nuclear RNA. The strongest AGO2 binding sites were mapped to micro RNAs (miRNAs). The most abundant miRNAs were distributed similarly between cytoplasm and nucleus, providing no evidence for mechanisms that facilitate localization of miRNAs in one compartment versus the other. Beyond miRNAs, most statistically-significant AGO2 binding was within introns. Splicing changes were confirmed by RT-PCR and were recapitulated by synthetic duplex RNAs and miRNA mimics complementary to the sites of AGO2 binding. These data support the hypothesis that miRNAs can control gene splicing. While nuclear RNAi proteins have the potential to be a natural regulatory mechanism, careful study will be necessary to identify critical RNA drivers of normal physiology and disease.

## INTRODUCTION

The power of RNA interference (RNAi) to repress translation in the cytoplasm of mammalian cells is well-known (Bartel, 2018). Argonaute (AGO) proteins are the primary protein factors that facilitate RNAi (Meister, 2013). There are four AGO proteins in human cells, AGO1-4. AGO2 is the best-studied and is the primary AGO variant capable of cleaving target RNAs (Liu et al., 2004; Meister et al. 2004). The roles of AGO1 and AGO3 are less clear while AGO4 has been observed to make the least contribution to RNAi (Petri et al., 2011). Recently, AGO3 has been reported to be catalytically activated by shorter guide RNAs (Park et al., 2020).

The first step of endogenous regulation of physiologic processes in the cytoplasm involves the recognition of microRNAs (miRNAs) by AGO proteins. The miRNA:AGO complex is generally assumed to recognize sequences within the 3’-untranslated region (3’-UTR) of genes through complementary binding to a “seed sequence” at bases 2-8 of the miRNA, leading to repression of translation.

While most attention has been focused on the impact of RNAi on translation (Zeng and Cullen, 2002), miRNAs and RNAi protein factors (including AGO variants) also reside in cell nuclei (Robb et al., 2005: Jeffries et al., 2011; Gagnon et al. 2014a). The presence of key components of the RNAi machinery in cell nuclei suggests that miRNAs may also have the potential to control transcription, splicing and other nuclear processes.

Evidence supporting the hypothesis that nuclear RNAi controls gene expression includes reports that small synthetic RNAs can be used to control transcription or splicing (Kalantari et al., 2016). Several laboratories have reported that promoter-target duplex RNAs can either repress or up-regulate transcription (Morris et al. 2004; Janowski et al. 2005; Janowski et al. 2007; Li et al., 2006; Huang et al., 2010; Matsui et al., 2010; Matsui et al. 2013; Zhang et al., 2014; Portnoy et al., 2016). Transcription can be controlled by synthetic miRNAs (Place et al., 2008; Kim et al., 2008; Younger and Corey 2011), and an endogenous miRNA can control endogenous cyclooxygenase 2 (COX-2) expression by binding to a transcript that overlaps the COX-2 promoter (Matsui et al., 2013).

Small RNAs can also regulate splicing (Allo et al., 2009; Liu et al., 2012). Small RNAs that target key regions near intron/exon boundaries have been shown to modulate splicing of an engineered luciferase model gene, dystrophin, and survival motor neuron 2 (SMN2) (Liu et al., 2012). This mechanism is like that of antisense oligonucleotides that target the same sites to affect alternative splicing by blocking binding of splicing factors (Havens and Hastings, 2016). Small synthetic RNAs can also target intronic RNA to couple chromatin silencing and alternative splicing (Allo et al., 2009; Ameyar-Zazoua et al., 2012). These studies imply that RNA:AGO complexes may be acting as ribonucleoprotein splicing factors, combining the versatile recognition properties of RNA with the stabilizing and bridging properties of proteins.

While these data related to the nuclear activities of synthetic duplex RNAs are intriguing, there have been no reports of endogenous miRNAs controlling splicing. An outstanding question, therefore, is whether the robust and versatile control achieved using designed synthetic RNAs reflects an unappreciated layer of natural regulation of transcription and splicing by small RNAs. To begin to answer this question it is necessary to understand where miRNA recognition occurs in the nuclear transcriptome and what consequences that recognition has for gene expression.

Here we use enhanced crosslinking immunoprecipitation (eCLIP) (Van Nostrand et al., 2016) to determine AGO2 localization within the transcriptome of mammalian cell nuclei. We then examined the effects of knocking out *AGO1*, *AGO2*, *AGO1/2*, *AGO1/2/3*, and *DROSHA* on alternative splicing for genes with significant intronic AGO2-binding sites. Our results suggest that AGO2 protein binds to sites with introns and that support the conclusion that alternative splicing can be achieved by miRNAs. TNRC6 proteins that bind AGO proteins and act in concert during RNAi exhibit a similar impact on alternative splicing (see accompanying manuscript, Johnson et al.)

## RESULTS

### Experimental design

We anticipated that it would be necessary to knock out more than one *AGO* variant. Therefore, we focused our analysis on HCT116 colorectal cancer-derived cells because they are diploid, facilitating the knockout of multiple genes. Another advantage is that miRNA expression in HCT116 cells is representative of miRNA expression as measured in a comprehensive study of approximately 1000 cancer cell lines (Ghandi et al. 2019), making it reasonable to expect that our data would be representative of many other widely used cultured cell lines. We also have *DROSHA* and *TNRC6* paralogs (see accompanying paper, Johnson et al.) knocked out in HCT116 cells, allowing direct comparison with other components of the RNAi pathway.

The AGO protein variants are functionally redundant in miRNA silencing (Su et al. 2009) suggesting that multiple knockouts are required to achieve clear effects. Therefore, we obtained *AGO1*, *AGO2*, *AGO1/2*, and *AGO1/2/3* knock out cells (**Figure 1AC**) (Chu et al., 2020). In HCT116 cells (Liu et al., 2018; Liu et al. 2019) and other cell lines (Petri et al., 2011), AGO4 protein is expressed at barely detectable levels and its knockout was not pursued.

**Figure 1.**
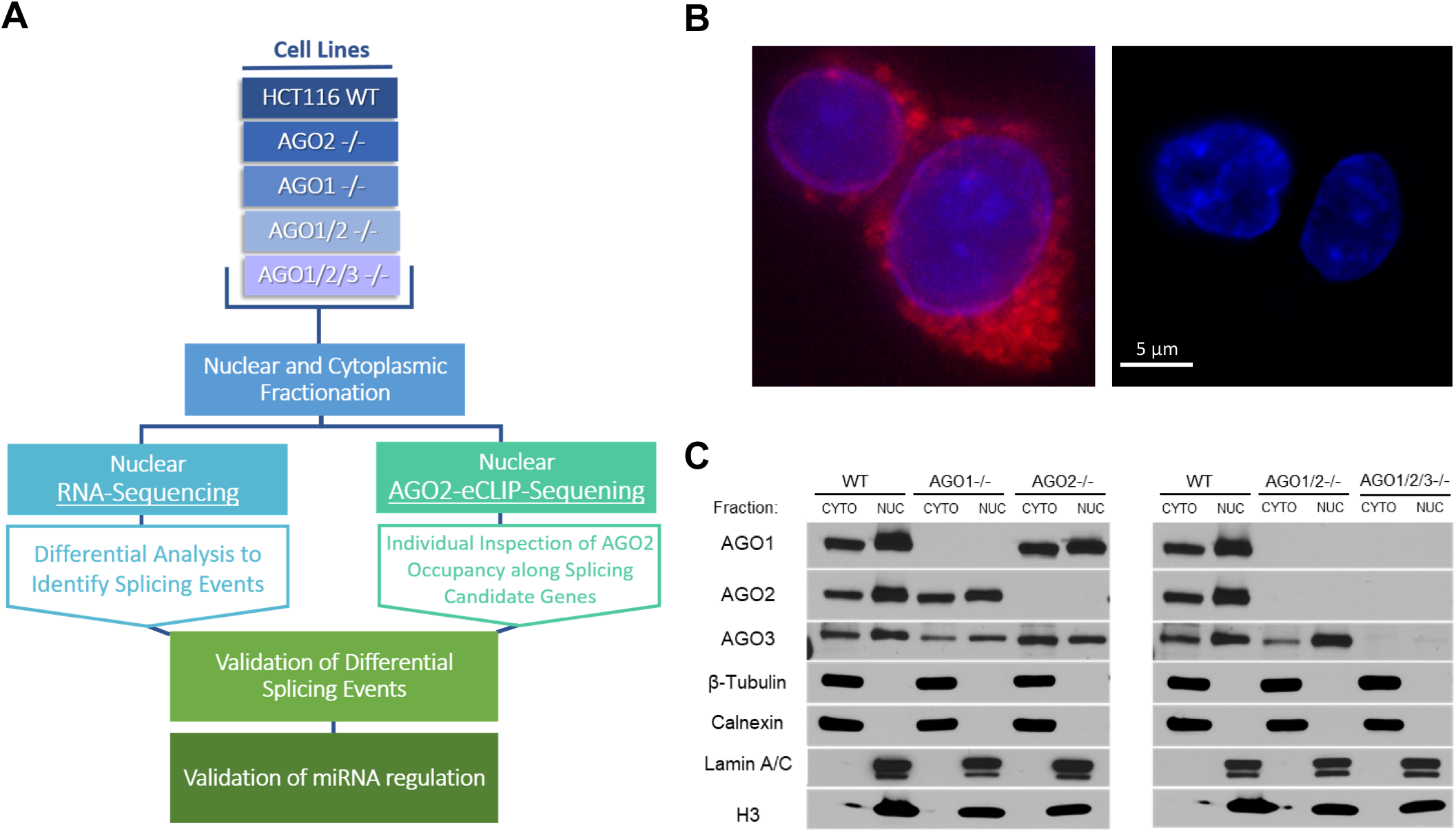
Experimental scheme for AGO2-eCLIP-sequencing. (*A*) Scheme showing knockout of *AGO* variants, eCLIP, and nuclear RNAseq analysis of alternative splicing relative to wild-type cells. (*B*) Microscopic imaging confirming the nuclei is free of ER contamination. (Left) Nuclei after one time wash with hypotonic buffer containing 0.5% NP40. (Right) Nuclei after 7 times wash with hypotonic buffer containing 2.0% NP40. blue: nuclear staining with DAPI; red: Endoplasmic reticulum marker. (*C*) Western blot validating the *AGO1*, *AGO2*, *AGO1/2*, and *AGO1/2/3* knockout cell lines and demonstrating nuclear fraction AGO protein levels and purity (representative of n=3).

We used purified nuclei for this study. Obtaining sufficiently pure nuclei is necessary because the endoplasmic reticulum contains AGO protein and is contiguous with the nuclear membrane (Stadler et al., 2013). The ER must be removed without disrupting the nuclei. For HCT116 cells, previous protocols (Gagnon et al., 2014a; Gagnon et al. 2014b) for removing ER from nuclei were not adequate for this cell line and required methodical optimization to identify more effective conditions. Microscopy (**Figure 1B**) and western analysis (**Figure 1C**) confirmed the removal of proteins associated with the endoplasmic reticulum (ER) and the presence in nuclei of AGO proteins.

Enhanced Crosslinking Immunoprecipitation (eCLIP) is a sensitive technique for identifying binding sites between RNA and protein that has been optimized to reduce the potential for artefactual background interactions (Van Nostrand et al., 2016). For our primary eCLIP analysis, we chose anti-AGO2 antibody 3148 that had been well-characterized as efficient for pull-down experiments (Grey et al., 2010; Boudreau et al., 2014) and that we had previously used for AGO2 eCLIP for cytoplasmic RNA. We identified AGO2 in the purified nuclei of crosslinked wild-type HCT116 cells and confirmed the absence of AGO2 protein in *AGO2* knockout cells. The nuclear samples yielded between ten and twenty-five million usable reads, sufficient to provide adequate coverage for thousands of genes (**Supplemental Figure S1**).

### Location of AGO2 within nuclear RNA

eCLIP data yields “clusters” of overlapping sequencing reads that identify potential sites for AGO protein binding within cellular RNA. We had previously reported eCLIP data from the cytoplasm of HCT116 cells revealing that AGO-binding clusters were primarily localized to the 3’-untranslated region (3’-UTR) (Chu et al., 2020). These data were consistent with standard assumptions about the mechanism of action for miRNAs, AGO protein, and RNAi in which cytoplasmic regulation is achieved by miRNA recognition at sequences within 3’-UTRs.

For nuclear RNA, however, almost 10-fold more AGO2 protein binding clusters were localized within intronic RNA than within the 3’-UTR (**Figure 2A**). Clusters were also detected within noncoding RNA, coding sequence RNA, and 5’-untranslated regions. Clusters were ranked by significance (p<0.05). Of the 200 most significant clusters, 129 were within sequences that encode miRNAs (**Figure 2B**). miRNAs are loaded directly onto AGO2, while mRNAs have a less direct association with AGO2. The direct contact between AGO2 and miRNAs, in combination with the relatively high expression of many miRNAs relative to mRNAs, may explain the prevalence of detecting strong miRNA:AGO associations.

**Figure 2.**
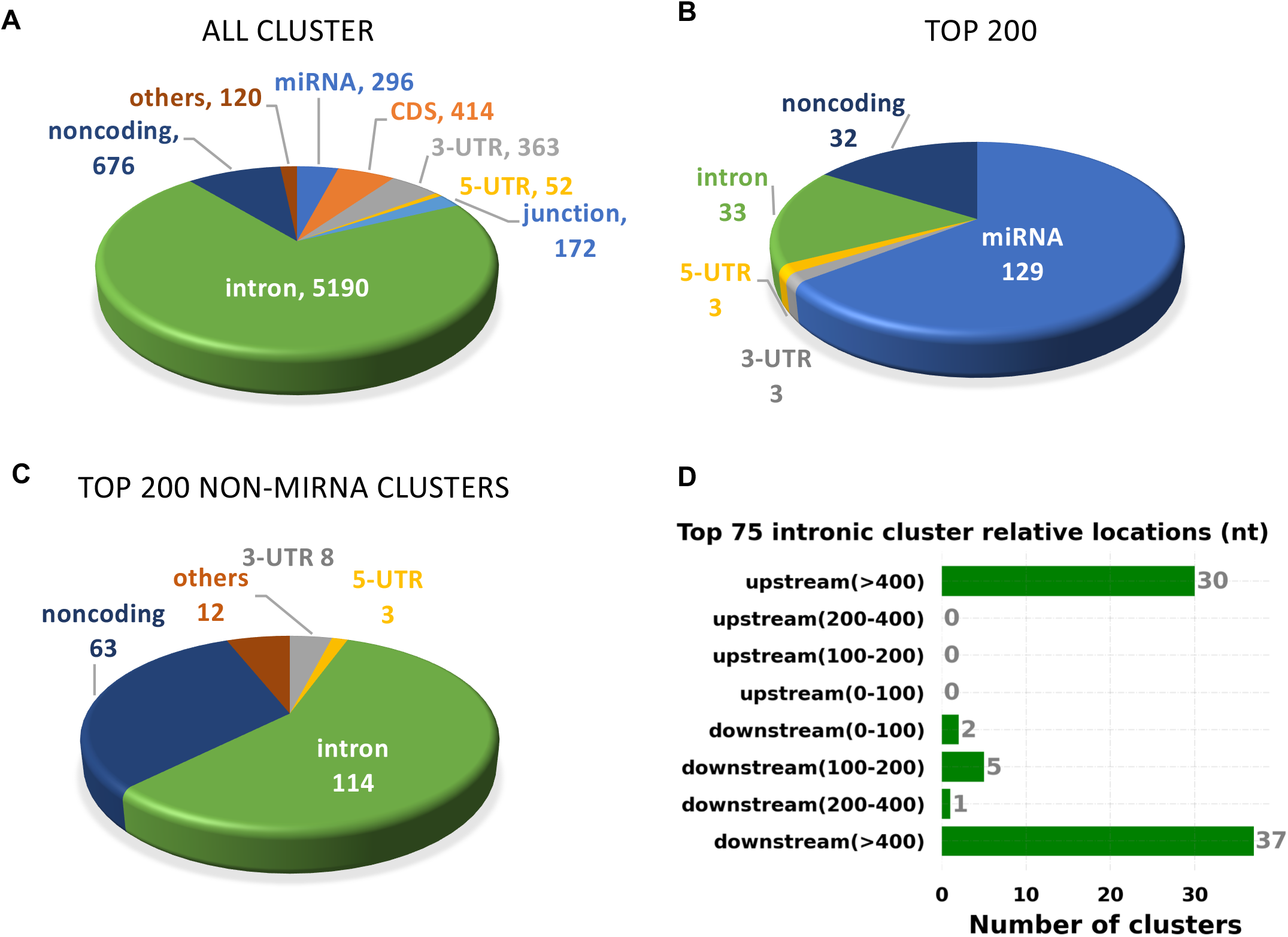
Localization of AGO2 protein engagement with nuclear RNA disclosed by eCLIP using an anti-AGO2. (*A*) Relative distribution of all AGO-binding clusters within nuclear RNA. (*B,C*) Relative distribution of top clusters, ranked by significance for the top 200 clusters overall and the top 200 clusters that that are not directly associated with miRNAs. (*D*) The top 75 AGO2 protein bound intronic cluster locations relative to the nearest splice sites. These 75 clusters were picked based on cluster *p*-value and enrichment fold over background.

To evaluate targets outside of sequences encoding miRNAs, we focused on the top 200 significant clusters that were not miRNAs (**Figure 2C**). 114 clusters were within introns and only eight within 3’-UTRs. The top-ranked intronic clusters tended to be distributed throughout intronic RNA rather than located near splice junctions (**Figure 2D**). The potential consequences of AGO2 binding within introns is discussed below.

### miRNA levels in cell nuclei mirror those in cell cytoplasm

Our laboratory and others have previously reported data from standard RNA-seq that show miRNAs are found in cell nuclei (Jeffries et al., 2011; Gagnon et al., 2014a). To further this analysis, we examined eCLIP-seq data for purified nuclear RNAs and compared that data to our previously published results evaluating eCLIP-seq data derived from cytoplasmic RNA (Chu et al., 2020).

We compared the number of eCLIP RNAseq reads for the sixty most expressed miRNAs in the nucleus to the read number of highly expressed miRNAs in the cytoplasm and observed similar trends in both cell compartments (**Figure 3A**). We also observed strong overlap among the one hundred miRNAs with highest read number (**Figure 3B**). The top eight miRNA families were the same in the nucleus and the cytoplasm (**Figure 3CD**). These highly ranked miRNAs accounting for ~ 60 % of all reads due to miRNAs. It is reasonable to expect that these highly ranked miRNAs will be most likely to have strong biological activities in HCT116 cells.

**Figure 3.**
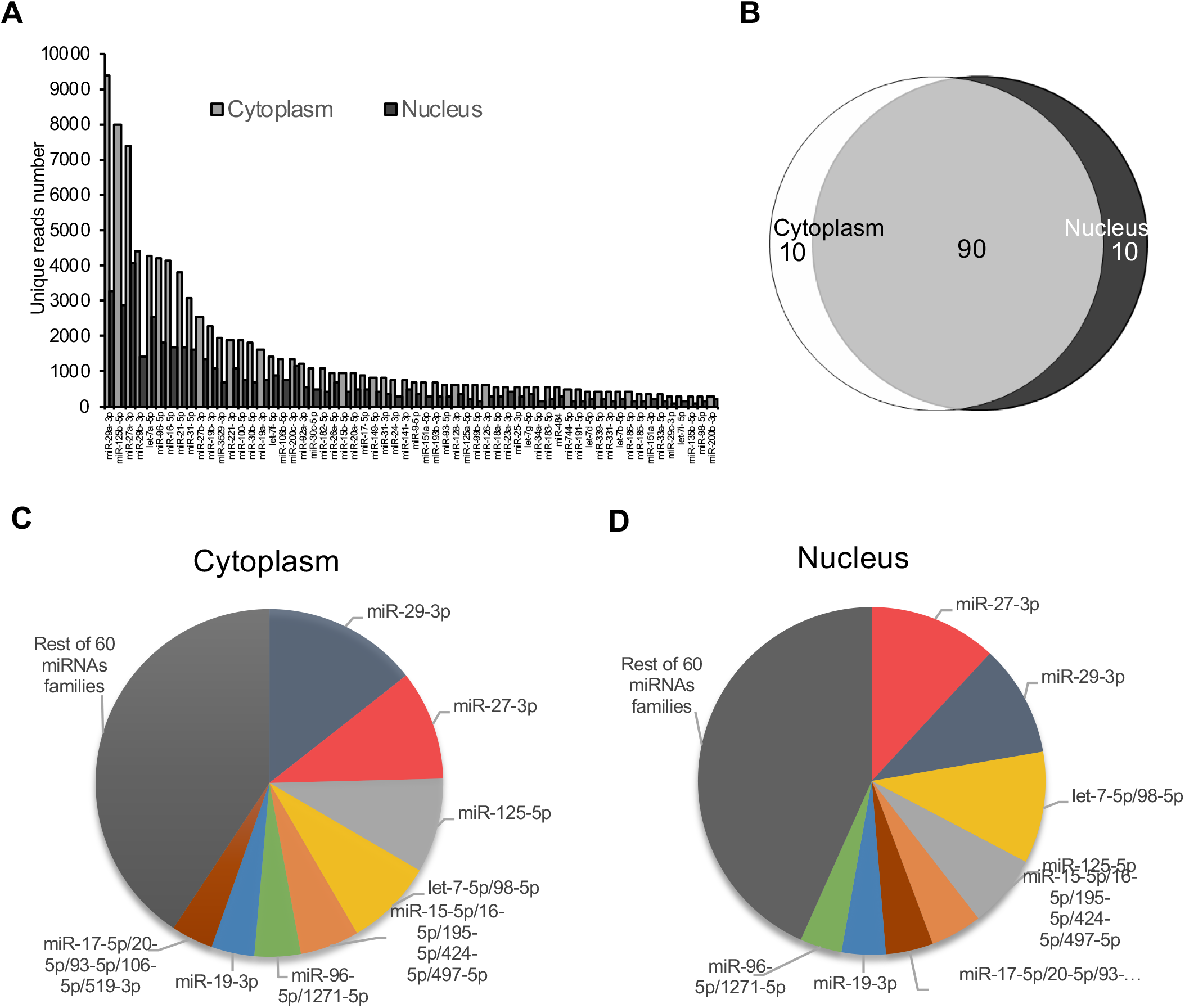
Rank ordering top AGO2-associated miRNAs in cytoplasm and nucleus. (*A*) miRNA abundance, ranked by the read number associated with each miRNA after eCLIP. (*B*) Overlap between top 100 miRNAs in nuclear and cytoplasmic fractions. (*C,D*) Percentage distribution of top one hundred miRNAs, grouped as families.

### Effect of AGO expression on alternative splicing

The predominance of AGO2 protein binding clusters within intronic RNA, in combination with previous observations that synthetic RNAs could target introns to alter splicing (Allo et al., 2009; Liu et al., 2012) led us to examine the hypothesis that AGO2 binding in concert with miRNAs might be an endogenous regulatory mechanism for controlling alternative splicing. We integrated our AGO2-eCLIP-seq and RNAseq data to identify genes that would be optimal candidates for detailed investigation of endogenous RNAi in the nucleus.

The impact on the total number of alternative splicing events was measured, including skipped exons, mutually exclusive exons, alternative 5’ splice sites, and alternative 3’ splice sites (**Figure 4A**). In the initial analysis, changes in the *AGO* knockout cell lines relative to wild-type cells were evaluated. We subsequently evaluated changes in splicing at genes with significant AGO2 binding clusters within introns relative to AGO knockout cells serving as negative controls or cells serving as size-matched input controls. We observed hundreds of changes in splicing (**Figure 4B**), but less than fifty changes were associated with AGO2 binding clusters within introns (**Figure 4C**).

**Figure 4.**
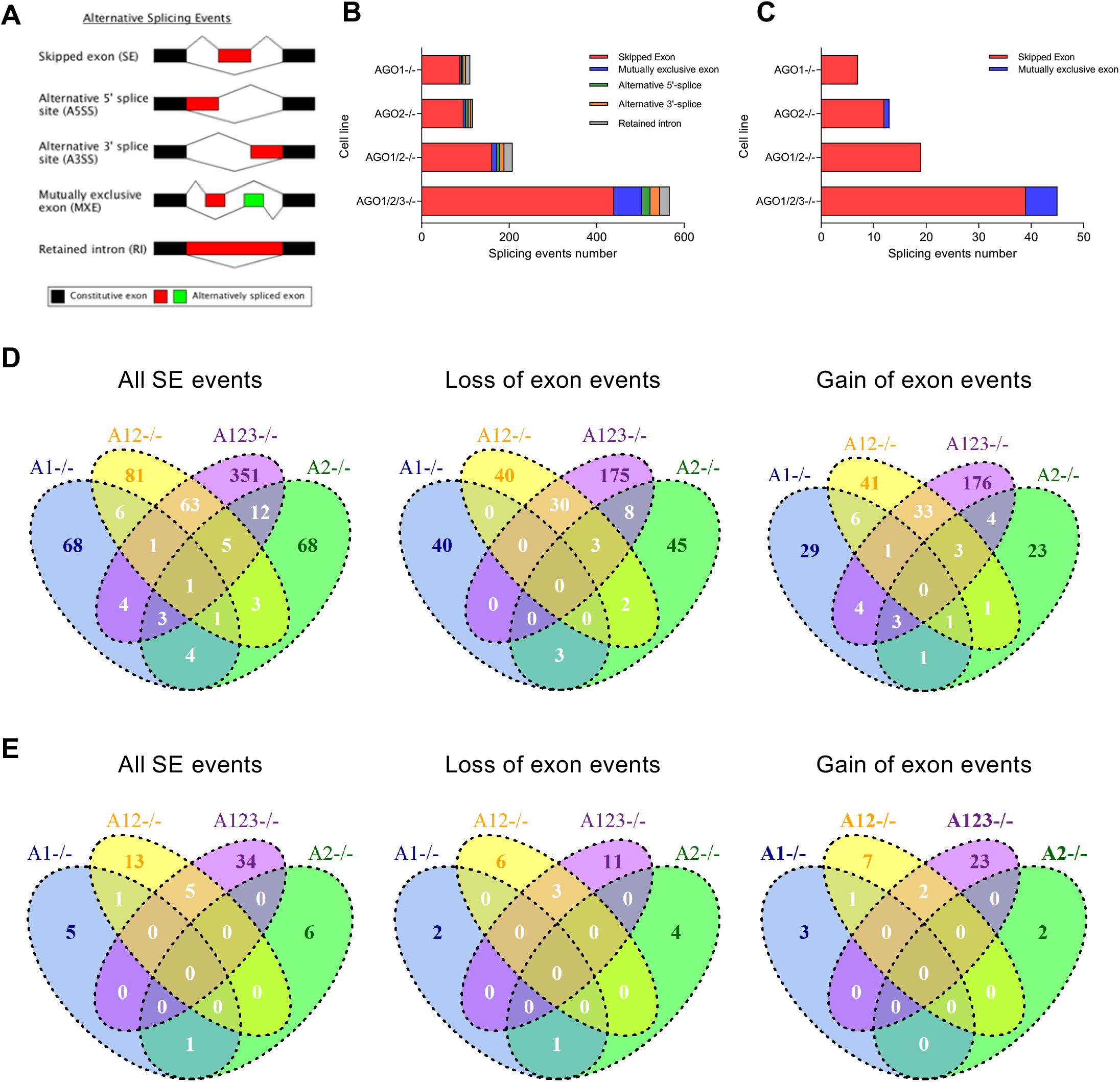
The effect of *AGO* knockouts on alternative splicing events. (*A*) Alternative splicing events. (*B*) The alternative splicing events observed upon *AGO* knockout. (*C*) Alternative splicing events associated with nearby AGO2 protein-binding sites. (*D*) VENN diagrams showing the overlap in skipped exon events for the various AGO knockout cell lines. (*E*) VENN diagrams showing the overlap of alternative splicing events for genes that have intronic binding sites for AGO2.

Experimental validation of results from large datasets is time-consuming and it is important to prioritize candidate genes for analysis. Some splicing events are likely to be indirect effects or noise unrelated to knocking out the *AGO* variants. We reasoned that the splicing events that are most likely to be endogenously regulated by RNAi would show significant AGO2 occupancy and be shared by more than on *AGO* knockout cell line, especially between the *AGO1/2* double knockout and the *AGO1/2/3* triple knockout.

We observed that, for loss or gain of exon events, several dozen significant events were shared between the *AGO1/2* and *AGO1/2/3* knockout cells (**Figure 4D**). For splicing events associated with location for AGO2 protein binding to introns and shared by *AGO1/2* and *AGO1/2/3* cells, we observed five skipped exon events, three loss of exon events, and two gain of exon events (**Figure 4E**). These splicing events identified through the combination of eCLIP and RNAseq are the strongest candidates for endogenous miRNA-mediated nuclear gene regulation.

### Experimental validation of splicing changes induced by AGO knockouts

We chose several splicing events for experimental validation based on their overlap in both *AGO1/2* and *AGO1/2/3* knockout cells (**Figure 4E**, **Supplemental Figures S2A-H**). These included exon exclusion events for *PHLDB1*, *FKBP14*, and *TBC1D5*, and exon inclusion events for *PPIP5K2*, *APIP*, *RUBCN*, and *KIF21A*. In all cases, we confirmed the alternative splicing predicted by our RNAseq data (**Figure 5**). Splicing changes were confirmed using a second PCR primer set for amplification (**Supplemental Figure S3**). For three genes, *PHLDB1*, *FKBP14*, and *APIP* the experimentally determined difference in splicing was observed in both the *AGO1/2* and *AGO1/2/3* knockout cells. For the other four genes, the difference in splicing was observed only in the triple knockout cells, but not the *AGO1/2* double knockout cells, possibly reflecting the impact of the remaining AGO3 protein.

**Figure 5.**
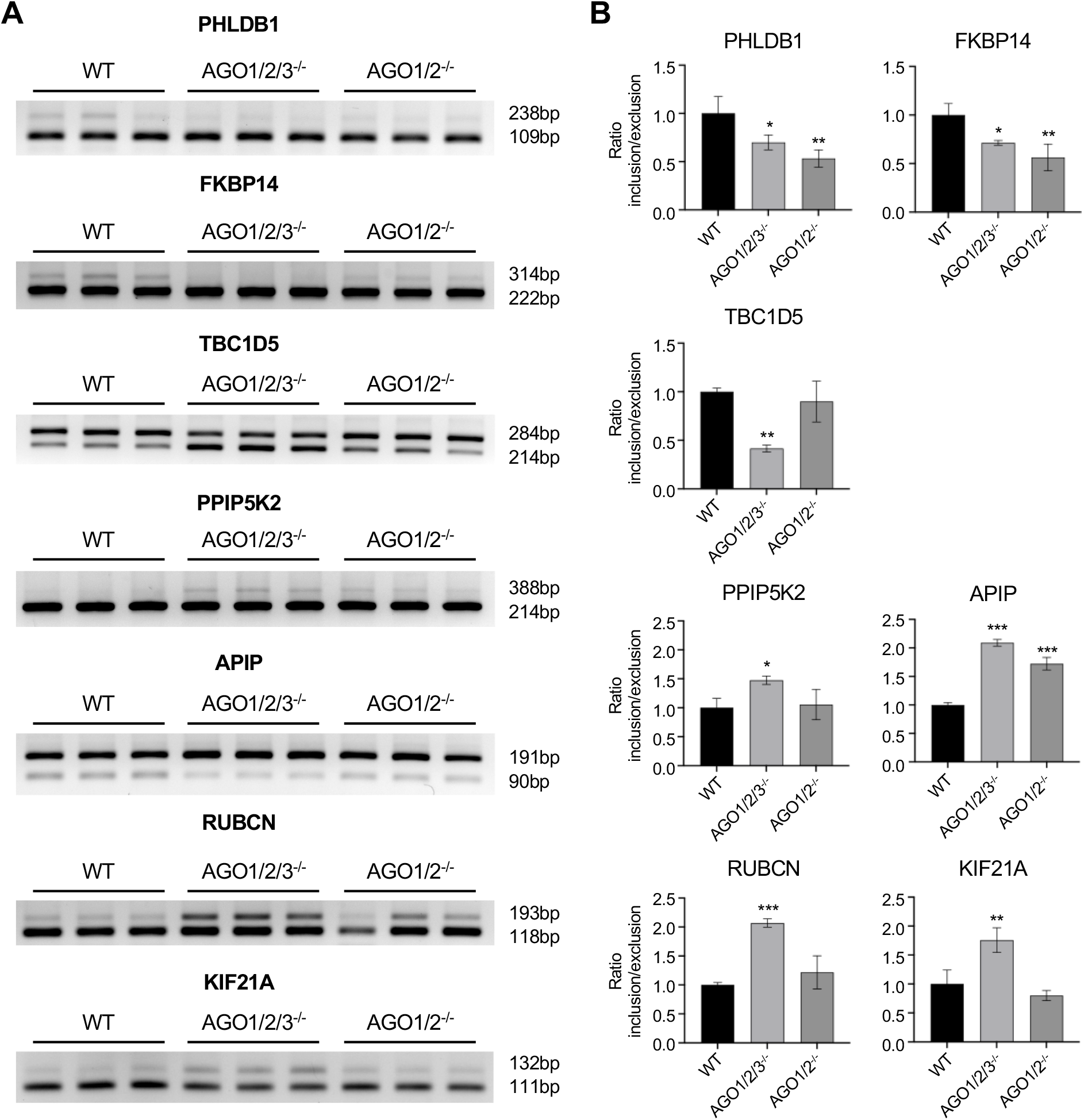
Validating the effect of *AGO* knockouts on alternative splicing. (*A*) Semiquantitative PCR validation of skipped exon events in *AGO1/2/3* KO and *AGO1/2 KO* cells. (*B*) Quantitation of data shown in in part (A). Error bars represent standard deviation (SD). *P < 0.05; **P < 0.01; ***P < 0.001 compared with WT by one-way ANOVA and Dunnett’s multiple comparisons test.

In an accompanying study, we examined the effect of knocking out *TNRC6*, a gene that produces a three paralog scaffolding proteins (TNRC6A, TNRC6B, and TNRC6C) that interact in concert with AGO2 protein. RNAseq from *TNRC6* knockout cells revealed many of the same splicing changes, and qPCR validation yielded splicing changes in *PHLDB1*, *FKBP14*, *RUBCN*, *KIF21A*, and *TBC1D5* (Johnson et al., 2021, accompanying paper).

### Effect of DROSHA knockout on splicing of select genes

DROSHA is an RNase III enzyme that catalyzes consecutive processing events during miRNA biogenesis (Michlewski and Caceres, 2019; Treiber et al., 2019). Like the AGO protein variants, DROSHA is an integral component of regulation by miRNAs. Knockout of DROSHA protein would be expected to have similar effects on gene expression and would tend to confirm the impact on splicing that we observe is due to impairing regulation by RNAi. We obtained *DROSHA* knockout cells (**Supplemental Figure S4**) and examined the impact of loss of *DROSHA* (**Figure 6**) on splicing of the seven genes chosen for experimental validation in our *AGO* knockout cells (**Figure 5**).

**Figure 6.**
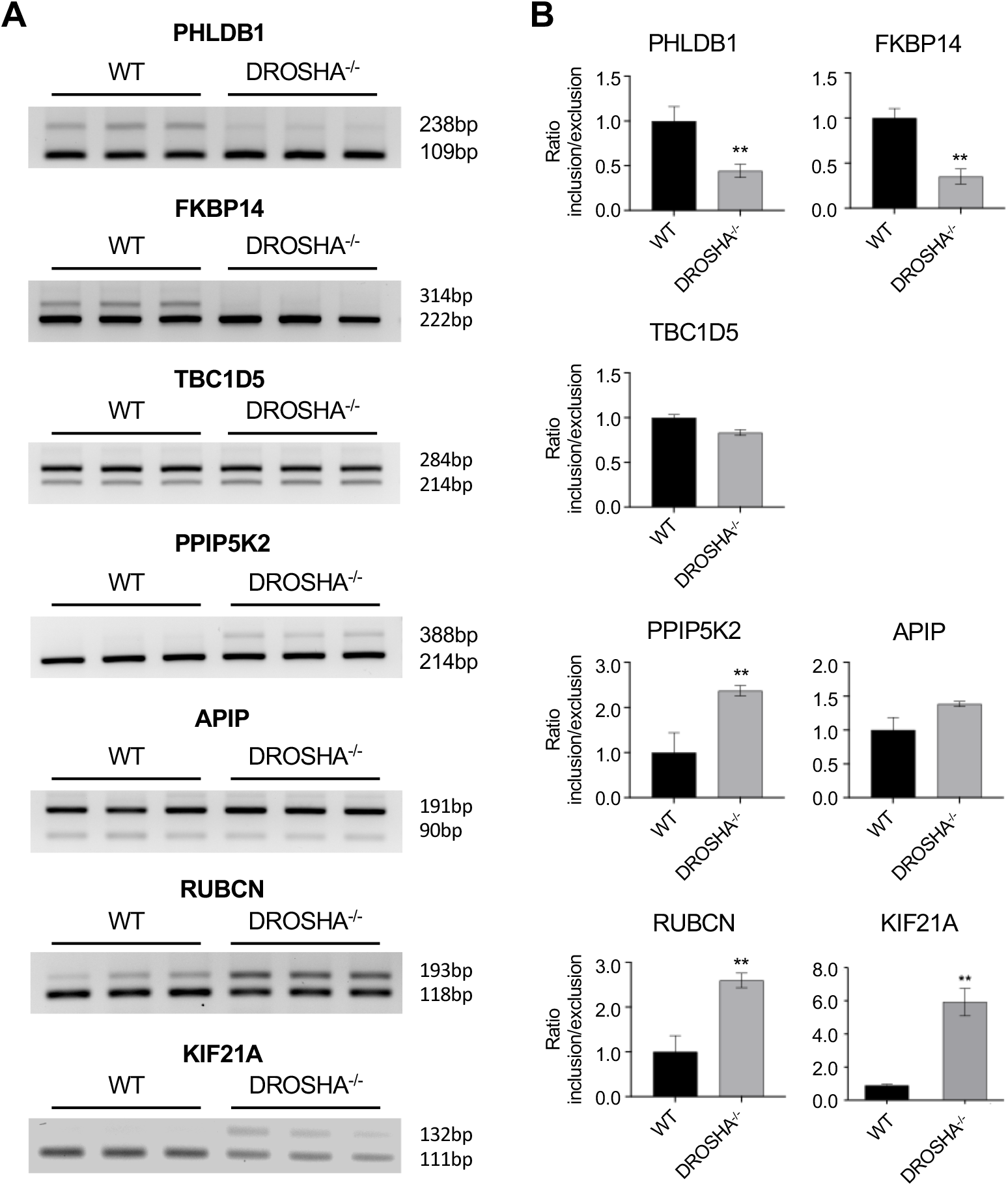
Impact of *DROSHA* knockout on skipped exon alternative splicing. Skipped exon events in *DROSHA* knockout cells. (*A*) Semiquantitative PCR validation of skipped exon events in *DROSHA* knockout cells. (*B*) Quantitation of data shown in in part (A). Error bars represent standard deviation (SD). *P < 0.05; **P < 0.01 compared with WT by t-test.

For five of the seven genes, we observed changes in alternative splicing consistent with our *AGO* knockout RNAseq data (**Figure 4**) and our experimental qPCR measurement of splicing changes after *AGO* knockout (**Figure 5**). We observed exon exclusion events for *PHLDB1* and *FKBP14* and exon inclusion events for *PPIP5K2*, *RUBCN*, and *KIF21A* (**Figure 6**). The alternative splicing changes for these five genes were confirmed using a second primer set for amplification (**Supplemental Figure S5**). Our similar data from loss of proteins from three different families (AGO, DROSHA, TNRC6) in the RNAi pathway support the hypothesis that the RNAi machinery can regulate alternative splicing.

### Alteration of splicing by synthetic miRNA mimics and anti-miRs

We used synthetic double-stranded RNAs and anti-miRs (**Table S1**) to test the link between RNA recognition within introns and regulation of alternative splicing. The synthetic RNAs were designed to target sequences with significant AGO protein binding clusters and that showed splicing changes in *DROSHA* and *AGO* knockout cells. We selected the sequences responsible for the AGO2 protein binding cluster in *RUBCN* and *FKBP14* intronic RNAs as targets for synthetic RNAs (**Figure 7AB**) because we had observed significant splicing changes in *AGO* (**Figures 5**), *DROSHA* (**Figure 6**), and *TNRC6* (see accompanying paper) knockout cells.

**Figure 7.**
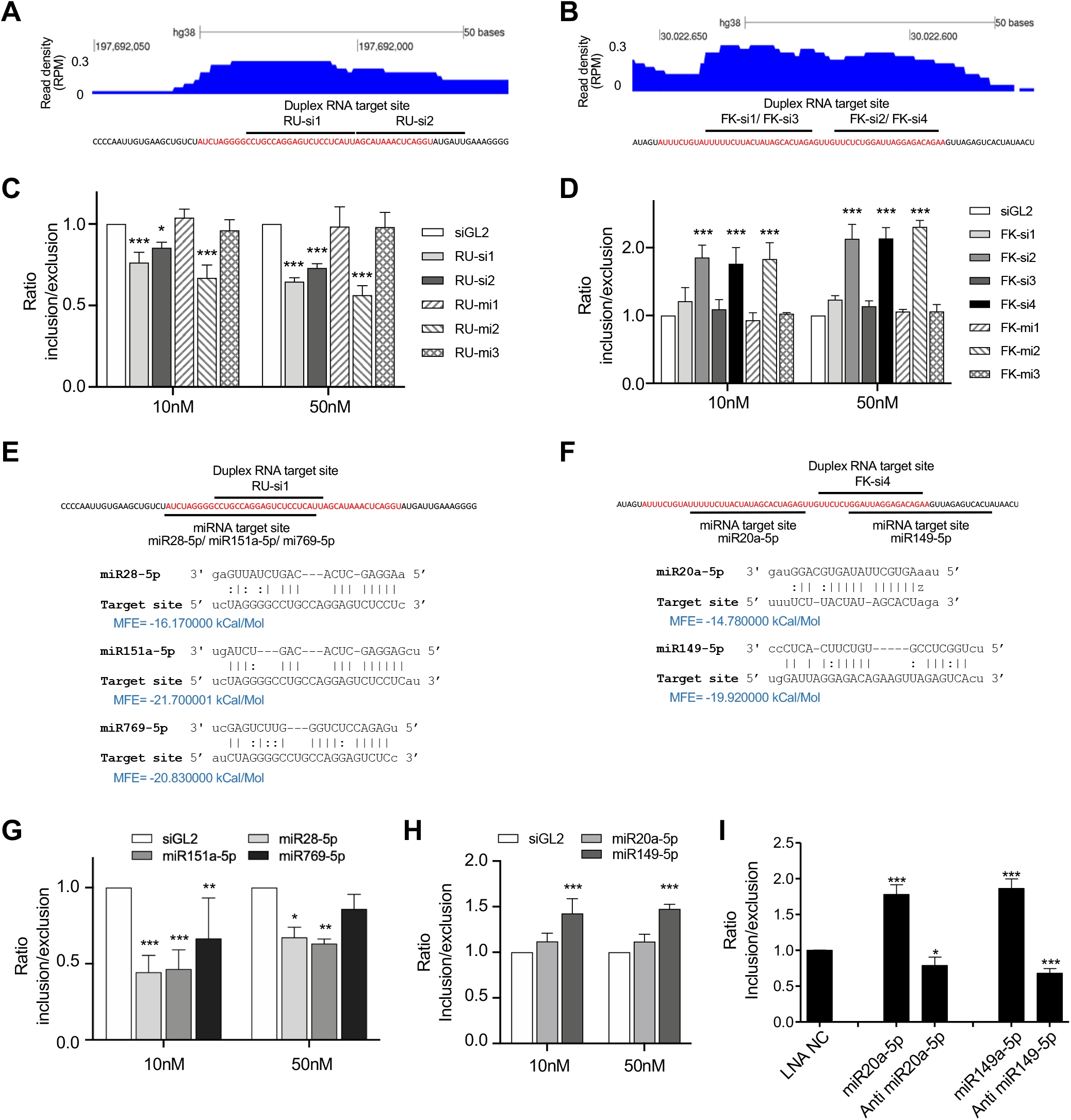
Double strand RNAs induce splicing changes in *RUBCN* and *FKBP14*. (*A*) AGO2 protein binding cluster within *RUBCN* intronic RNA. (*B*) AGO2 cluster within *FKBP14* intronic RNA. Red colored sequence is shown the significant AGO2 cluster analyzing from eCLIP-seq. (*C,D*) qPCR quantification of splicing changes by double strand RNAs in (*C*) *RUBCN* and (*D*) F*KBP14*. (*E,F*) Potential binding sites for miRNAs at AGO2-clusters within (*E*) *RUBCN* and (*F*) *FKBP14*. (*G,H*) qPCR quantification of splicing changes by miRNA mimics in (*G*) *RUBCN* and (*H*) *FKBP14*. (*I*) qPCR quantification of splicing changes by miRNA mimics in *FKBP14*. Error bars represent standard deviation (SD). *P < 0.05; **P < 0.01; ***P < 0.001 compared with WT by one-way ANOVA and Dunnett’s multiple comparisons test. siGL2 is endogenous non-targeting negative control siRNA. LNA-NC is non-targeting negative control ASO.

In these experiments, the ability of AGO2 protein to cleave RNA during RNAi is not necessary. Instead, simply blocking an important RNA sequence may be sufficient to alter splicing. Approaches that block RNA rather than cut it avoid cleavage of the introns that would confound our analysis. To avoid any potential for cleavage, we designed duplex RNAs to contain mismatches at positions 9 or 9/10 based on crystallographic analysis demonstrating that such mismatches disrupt catalysis (Wang et al., 2008).

When these duplex RNAs were transfected into HCT116 cells, we observed decreased exon inclusion (**Figure 7C**) when targeting *RUBCN* and increased exon inclusion (**Figure 7D**) when targeting *FKBP14*. Knocking out AGO (**Figure 5**), DROSHA (**Figure 6**), or TNRC6 protein (see accompanying paper) protein had the reverse effect, increased exon inclusion for *RUBCN* and decreased exon inclusion for *FKBP14*. The results are consistent with the intronic target site being susceptible to regulation by duplex RNAs.

Next, we examined those splicing clusters at the *RUBCN* and *FKBP14* genes for complementarity to miRNAs (**Figure 7EF**). To predict miRNAs that might bind at those AGO2 clusters, two criteria were set: 1) miRNAs were limited to the top 100 most highly ranked miRNAs from our RNAseq analysis; and 2) the minimum free energy (MFE) for matching at the site was required < −14 Kcal/mol. From this list of miRNA candidates, we focused on miRNAs that overlapped intronic AGO2-binding clusters that showed significant splicing changes. We identified three miRNAs for *RUBCN* (miR28-5p, miR151a-5p, miR769-5p) and two for *FKBP14* (miR20a-5p, miR149-5p). As we had observed for the designed synthetic RNAs, we identified miRNA mimics (**Table S2**) that decreased exon inclusion when targeting *RUBCN* and increased exon inclusion when targeting *FKBP14* (**Figure 7GH, Supplemental Figure S6**).

Anti-miRs are synthetic oligonucleotides that are complementary to miRNAs and have the potential to block their effects (Krutzfeldt et al., 2005). We introduced locked nucleic acid (LNA) anti-miRs into HCT116 cells that were designed to target two miRNAs, miR-20 and miR-149 with complementarity to the AGO cluster within FKBP14 intronic RNA. Both anti-miRs reduce exon inclusion, while addition of miRNA mimics in parallel yield the opposite effect (**Figure 7I, Supplemental Figure S6**).

## DISCUSSION

### Localizing AGO2 binding within the nuclear transcriptome

We had previously employed CLIP-seq (Chi et al., 2009; Hafner et al., 2010) to identify AGO protein binding sites but were not able to unambiguously validate any expression changes (results not shown). We found that roadblocks to successful application of CLIP-seq in these prior experiments included: 1) We were comparing wild-type cells to cells where AGO2 expression had been reduced by siRNA-mediated knockdown. Since the knockdown was not complete, residual AGO2 protein might confound our ability to interpret data: 2) The expression of the other AGO variants were not reduced, keeping a pool of AGO available for controlling gene expression; 3) While CLIP-seq is a powerful approach, artefactual identification of AGO binding clusters can occur. Experimentally validating clusters is time-consuming, and we found that discriminating among clusters to identify the most promising candidates for analysis was inefficient. For a novel cellular process like small RNA mediated control of splicing, a more defined experimental strategy was necessary.

To better discriminate against background in both CLIP and RNAseq experiments, we obtained *AGO2* knockout cells (Golden et al., 2017; Chu et al., 2020). We also obtained *AGO1/2* and *AGO1/2/3* knockout cells (Chu et al., 2020) to remove the three most highly expressed AGO protein variants. Finally, we used an improved CLIP-seq technique, eCLIP, to reduce background signal (Van Nostrand et al., 2016).

### Nuclear Localization of miRNAs

Our eCLIP data revealed that the strongest and most significant AGO2 protein binding clusters within nuclear RNA were associated with miRNAs (**Figure 2**). Strong association with miRNAs is probably due to the AGO:miRNA interaction being direct instead of the secondary interaction formed between AGO and mRNA.

Our data demonstrate that AGO2-bound miRNAs in HCT116 cell nuclei have abundances like those in cell cytoplasm. There is no evidence in HCT116 cells of dramatic differences in miRNA distribution, suggesting that there is no need to hypothesize mechanisms to preferentially shuttle select individual miRNAs to nuclei relative to cytoplasm or vice versa in this context. The presence of RNAi factors (Robb et al. 2005; Gagnon et al., 2014a) and miRNAs (at a similar abundance) in both cytoplasm and nuclei supports the conclusion that the RNAi has the potential to recognize nuclear RNA and control gene expression and that there may be similar rules governing the recognition of target sequences.

Our data from one cell line grown under standard conditions does not rule out the possibility that a miRNA might be preferentially localized to nucleus relative to cytoplasm or vice versa. However, the data do suggest that strong evidence should be shown to support the shuttling of individual miRNAs in future studies.

### Impact of AGO expression on alternative splicing

After miRNAs, the strongest AGO2 protein association was within intronic RNA (**Figure 2**). We and others had previously shown that synthetic duplex RNAs complementary to introns could be used to control splicing (Allo et al., 2009; Liu et al., 2011), leading to the hypothesis that endogenous miRNAs might also be involved in the control of alternative splicing. The obvious mechanism for controlling splicing would be for the AGO:miRNA complex to block recognition of splicing factors. This mechanism is widely used for antisense oligonucleotides that modulate splicing (Havens and Hastings, 2016) and we had previously observed that duplex RNAs that mimic the sequence of splice-modulating antisense oligonucleotides and affect alternative splicing (Liu et al., 2012).

We now report several observations that support the hypothesis that small RNAs can control splicing. When *AGO* expression is knocked out, we observe several hundred changes in alternative splicing. Dozens of these changes occur at genes that have significant clusters of sequencing reads overlapping nearby introns. We further prioritized candidates by requiring that they be observed in both *AGO1/2* and *AGO1/2/3* knockout cells.

The significant splicing changes that were detected by RNAseq were experimentally validated by RT-PCR. Those same genes showed similar changes in splicing upon knockout of DROSHA, a processing enzyme that is an upstream component of the RNAi machinery. TNRC6 is a critical RNAi scaffolding protein that binds to AGO and several of these top candidate genes also showed splicing changes in TNRC6 knockout cells (see accompanying manuscript). Finally, both duplex RNAs, miRNA mimics, and miRNA inhibitors (anti-miRs) that target potential miRNA binding sites within introns were shown to regulate alternative splicing. Taken together, these data support the conclusion that miRNAs and the endogenous cellular RNAi machinery can regulate alternative splicing in HCT116 cells.

However, while our data suggest that duplex RNAs can regulate alternative splicing, the number of genes affected was modest. Changes in only seventy alternative splicing events are observed in both *AGO1/2* and *AGO1/2/3* knockout cells. For alternative splicing events that are associated with significant AGO binding clusters and that are observed in both *AGO1/2* and *AGO1/2/3* knockout cells, we observe just five skipped exon events, three loss of exon events, and two gain of exon events. A similar small number of events is observed in *TNRC6* knockout cells (see accompanying manuscript).

Why were so few events detected? We do not claim that we have identified all alternative splicing events that are candidates for regulation by miRNAs. For example, intronic RNA that is present at low copy numbers might be biologically significant but remain undetected during eCLIP. Our criteria for selecting candidates (significantly altered splicing in both *AGO1/2/3* and *AGO1/2/3* knockout cells for genes with significant intronic AGO2 protein binding clusters) was stringent and might also have led us to miss biologically relevant alternative splicing events.

Our HCT116 cells were grown under standard, permissive cell culture conditions. It is also possible that that under standard cell culture conditions the impact of the cellular RNA machinery and the number of RNA-regulated splicing events is small. miRNAs may exert their most robust effects under a limited range of environmental or growth conditions. miRNAs that control splicing would be no exception to this model – a potentially powerful regulatory mechanism that becomes important at critical points during development, response to environment change, or disease pathogenesis. If true, the modest changes in splicing that we observe in HCT116 cells might hint at the possibility that a larger range of changes occurs in other settings.

### Conclusions

AGO:miRNA complexes are ribonucleoproteins that have the potential to recognize sequences throughout the transcriptome. This ability coupled with the presence of both AGO proteins and miRNAs in the nucleus suggests the potential to control gene splicing. Consistent with this hypothesis we observe AGO2 binding within intronic RNA and identify endogenous miRNAs that may affect alternative splicing. Our data expand the potential for RNAi to control gene expression and miRNAs that affect splicing may play significant roles in physiology and disease.

## Materials and Methods

### Cell lines

The HCT116 cell line (Horizon Discovery) originated from the American Type Culture Collection (ATCC). ATCC authenticated this HCT116 cell line using Short Tandem Repeat (STR) analysis as described (Capes-Davis). The ATCC STR analysis compared seventeen short tandem repeat loci plus the gender determining locus, Amelogenin, to verify the HCT116 cell line (ATCC CCL 247). The European Collection of Authenticated Cell Cultures (ECACC) performed an additional STR analysis of seventeen loci on the cells received from ATCC, and the verified HCT116 cells (ECACC 91091005) were supplied to Horizon Discovery for distribution.

For eCLIP, we used HCT116:AGO2 knockout cells obtained from Joshua T. Mendell (UT Southwestern). The *AGO1*, *AGO2*, *AGO1/2*, and *AGO1/2/3* knockout cell lines used for RNAseq were prepared using GenCRISPR™ gene editing technology and services (GenScript) and verified HCT116 cells (Horizon Discovery). The AGO2 knockout cell line was independently generated to ensure that all cell lines used for RNAseq were derived from similar genetic backgrounds. gRNA sequence, gRNA target locations, and DNA sequencing results have been reported previously (Chu et al., 2020). The *DROSHA* knockout cell lines used were prepared using GenCRISPR™ gene editing technology and services (GenScript) and verified HCT116 cells (Horizon Discovery). gRNA sequence, gRNA target locations, and DNA sequencing results were shown in **Supplemental Figure S4**. All HCT116 and HCT116-derived cells were cultured in McCoy’s 5A Medium (Sigma-Aldrich) supplemented with final 10% FBS at 37 °C in 5% CO_2_. For the cell growth assay, the cells were seeded at a density of 50,000 cells/mL, disassociated with 1X trypsin, and counted using trypan blue staining (TC20 Automated Cell Counter, Bio-Rad).

### Preparation of nuclear extract

Nuclear extract isolation was similar as previously described (Gagnon et al., 2014a; Gagnon et al. 2014b), although modifications to adapt the protocol for HCT116 and HCT116-derived cells were required. Cells at ~95% confluence were lysed in hypotonic lysis buffer (HLB) (10 mM Tris-HCl, pH-7.4, 10 mM NaCl, 3 mM MgCl_2_, 2.5% NP-40, 0.5 mM DTT, 1x protease inhibitor (Roche) and 50 U/mL ribonuclease inhibitors (RNasin^®^ Plus, Promega) and supernatant collected as cytoplasmic fraction. Western blots to determine the purity of fractions and RNAi factors distribution were performed using western-blot analysis as before (Gagnon et al. 2014ab).

### Western-blot analysis

Total protein lysate was prepared re-suspending cells in lysis buffer (50 mM Tris-HCl, pH-7.0, 120 mM NaCl, 0.5% NP-40, 1 mM EDTA, 1 mM DTT, 1x protease inhibitor (Roche, cOmplete). Protein were separated on 4-20% gradient Mini-PROTEAN^®^ TGXTM precast gels (Bio-Rad). After gel electrophoresis, proteins were wet transferred to nitrocellulose membrane (0.45 μm, GE Healthcare Life Sciences) at 100 V for 75 min. Membranes were blocked for 1 hour at room temperature with 5% milk in 1x PBS containing 0.05% TWEEN-20. Blocked membranes were incubated with the primary antibodies in blocking buffer at 4 °C on rocking platform overnight: using anti-AGO1, 1:2000 (5053, Cell Signaling), anti-AGO2, 1:1500 (015-22031, Fujifilm WAKO), anti-AGO3, 1:500 (39787, Active Motif), anti-Calnexin, 1:1000 (2433, Cell Signaling), anti-LaminA/C, 1:1500 (ab8984, Abcam), anti-β-Tubulin, 1:5000 (T5201, Sigma-Aldrich), anti-Histone H3, 1:20000 (2650S, Cell Signaling) antibodies. After primary antibody incubation, membranes were washed 3 x 10 min at room temperature with 1xPBS+0.05% TWEEN-20 (PBST 0.05%) and then incubated for one hour at room temperature with respective secondary antibodies in blocking buffer. Membranes were washed again 4 x 10 min in PBST 0.05%. Washed membranes were soaked with HRP substrate according to manufacturer’s recommendations (SuperSignal™ West Pico Chemiluminescent substrate, Thermo Scientific) and exposed to films. The films were scanned, and bands were quantified using ImageJ software.

### Cell Transfection

For experiments of double strand RNAs and miRNA mimics, cells were plated onto 48-well plates at 30,000 cells per well 1 days prior to transfection. Cells were transfected with double strand RNAs or miRNA mimics by using Lipofectamine 3000 (Invitrogen). Detailed sequence information is shown in Supplementary Table S4. miRCURY LNATM miRNA inhibitor for anti-miR20a-5p and anti-miR149-5p were bought from Qiagen. Total RNAs were extracted 24 hours or 96 hours (for anti-miR test) after transfection with TRIzol (Invitrogen) for RT-PCR.

### Splicing analysis by gel electrophoresis and qPCR

Total RNA was extracted from HCT116 wild-type, knockout cells, and transfected cells, and treated with DNase I (Worthington Biochemical) at 25 °C for 20 min, 75 °C for 10 min. Reverse transcription was performed using high-capacity reverse transcription kit (Applied Biosystems) per the manufacturer’s protocol. 2.0 μg of total RNA was used per 20 μl of reaction mixture. In gel electrophoresis analysis, PCR amplification was as follow; 95 °C 5 min and 95 °C 15 s, 60 °C 1 min for 38 cycles. The PCR products were separated by 1.5% agarose gel electrophoresis. The bands were quantified by using ImageJ software. In qPCR analysis for splicing changes by using double strand RNAs and miRNA mimics, PCR was performed on a 7500 real-time PCR system (Applied Biosystems) using iTaq SYBR Green Supermix (BioRad). PCR reactions were done in triplicates at 55 °C 2 min, 95 °C 3 min and 95 °C 20 s, 60 °C 1 min for 40 cycles in an optical 96-well plate. The expression level was compared between exon inclusion variants and exon exclusion variants. PCR primers were shown in **Supplementary Table S2**.

### RNAseq for gene expression analysis

WT HCT116, AGO1, AGO2, AGO1/2, and AGO1/2/3 knock out cells were used for RNAseq. Three biological replicated samples were sequenced. Approximately 3.0×10^6^ cells were seeded in a 15-cm large dish. Cells were harvested 48 hours later and RNA was extracted from cytoplasmic or nuclear fractions using the RNeasy Mini Kit (Qiagen) with an on-column DNase digestion. Sequencing libraries were generated using the TruSeq Stranded Total RNA with Ribo-Zero Human/ Mouse/Rat Low-throughput (LT) kit (Illumina) and run on a NextSeq 500 for paired-end sequencing using the NextSeq 500/550 High Output v2 Kit, 150 cycles (Illumina).

Quality assessment of the RNAseq data was done using NGS-QC-Toolkit43 with default settings. Quality-filtered reads generated by the tool were then aligned to the human reference genome hg38 and transcriptome gencode v75 using the STAR (v 2.5.2b) using default settings. Read counts obtained from STAR were used as input for Salmon (v 1.0.0) and Deseq2 for gene differential expression analysis. Genes with adjusted *p* ≤ 0.05 were regarded as differentially expressed for comparisons of each sample group.

### eCLIP

Control and AGO2 ^−/−^ HCT116 cells (obtained from Dr. Joshua Mendel, UT Southwestern) were seeded in 15 cm dishes with 12 dishes per cell line at 3.0×10^6^ cells per dish. Cells were cultured for 48 hours and subsequently UV crosslinked at 300 mJ/cm^2^. Nuclear fraction was collected as described above. eCLIP was performed using the frozen samples as previously described (Van Nostrand et al., 2016), using anti-AGO2 antibody for IPs (3148, gift from Jay A. Nelson lab). For each cell line, duplicate input and IP samples were prepared and sequenced. The RiL19 RNA adapter (**Supplementary Table S3**) was used as the 3’ RNA linker for input samples. RNA adapters RNA_A01, RNA_B06, RNA_C01, RNA_D08, RNA_X1A, RNA_X1B, RNA_X2A, RNA_X2B were used for IP samples (**Supplementary Table S3**). PAGE purified DNA oligonucleotides were obtained from IDT for the PCR library amplification step (**Supplementary Table S3**). PCR amplification was performed using between 11-16 cycles for all samples. Paired-end sequencing was performed on a NextSeq 500 using the NextSeq 500/550 High Output v2 Kit, 100 cycle (Illumina). eCLIP data was analyzed as previously described (Chu et al. 2020). Initial AGO2 binding clusters identified by CLIPper (Lovci et al., 2013) in wild-type HCT116 were filtered to keep only clusters that are statistically significant (p < 0.001). For each region, normalization to total usable reads was performed and a fold change between IP and combined samples (input and IP in knockout cell line samples) was calculated. Significant CLIP clusters in each dataset were defined by i) *p* < 0.05 determined by the Fisher exact test or Yates’ Chi-Square test, and ii) log2 fold change of normalized reads in the cluster was ≥ 2 comparing IP to combined (input + IP in knockout cells).

The final CLIP clusters for AGO2 were identified by first identifying significant clusters present in both experimental replicates. A cluster was considered to be present in both replicates if it occurred on the same strand and the replicate clusters overlapped by at least 1/3 of their total length. Significant clusters from both replicates were then merged to define the final cluster length. Clusters were annotated based on their genomic locations (Gencode v27). If a cluster was assigned to multiple annotations, the annotation was selected using the following priority: CDS exon > 3’ UTR > 5’ UTR > Protein-coding gene intron > Noncoding RNA exon > Noncoding RNA intron > Intergenic.

### Estimation of miRNAs Expression Levels from eCLIP-seq

Bowtie2 was used to map the miRNA tags to the miRBase (mature miRNAs), allowing up to one mismatch. The miRNA expression levels are quantified as the number of reads mapped to individual miRNAs normalized by the total number of mapped reads in miRBase. The expression levels from different samples were further normalized by quantile normalization to control for batch effect.

### Statistical analysis

The dynamic and bar graphs represent mean and standard deviation. The averages among cells we compared using one or two ways analysis of variance followed by Bonferroni and Tukey post-hoc tests (p<0.05). To determine correlation between AGO2 binding cluster significance level and gene expression change in AGO knockout cell, we first tested data sets for normal distribution (D’Agostino and Pearson omnibus normality test, Kolmogorov-Sminov test). Both correlating data sets could not pass normality test, or in many cases relation was not linear, therefore we calculated Spearman’s correlation coefficient.

### Data availability

All high-throughput sequencing data generated for this study (RNAseq, eCLIP) have been deposited in Gene Expression Omnibus under accession number GSE161559.

## SUPPLEMENTAL MATERIAL

Supplemental material is available for this article.

## ACKNOWLEDGEMENTS

We thank Dr. Bethany Janowski for critical reading of this manuscript. The authors thank Dr. Joshua Mendell for the gift of HCT116:AGO2 knock out cells and Dr. Jay Nelson for anti-AGO antibody 3148. DRC was supported by the National Institutes of Health (NIH) (GM106151) and the Robert Welch Foundation (I-1244). DRC holds the Rusty Kelley Professorship in Medical Science. KCJ was supported by a Fellowship from the NIH (1F31GM137591-01).

## Author contributions

D.R.C. wrote the manuscript and supervised the experiments. Y.C., A.K., K.C.J., S.Y., and J.L. performed the experiments.

---

**Supplemental Figure S1.**
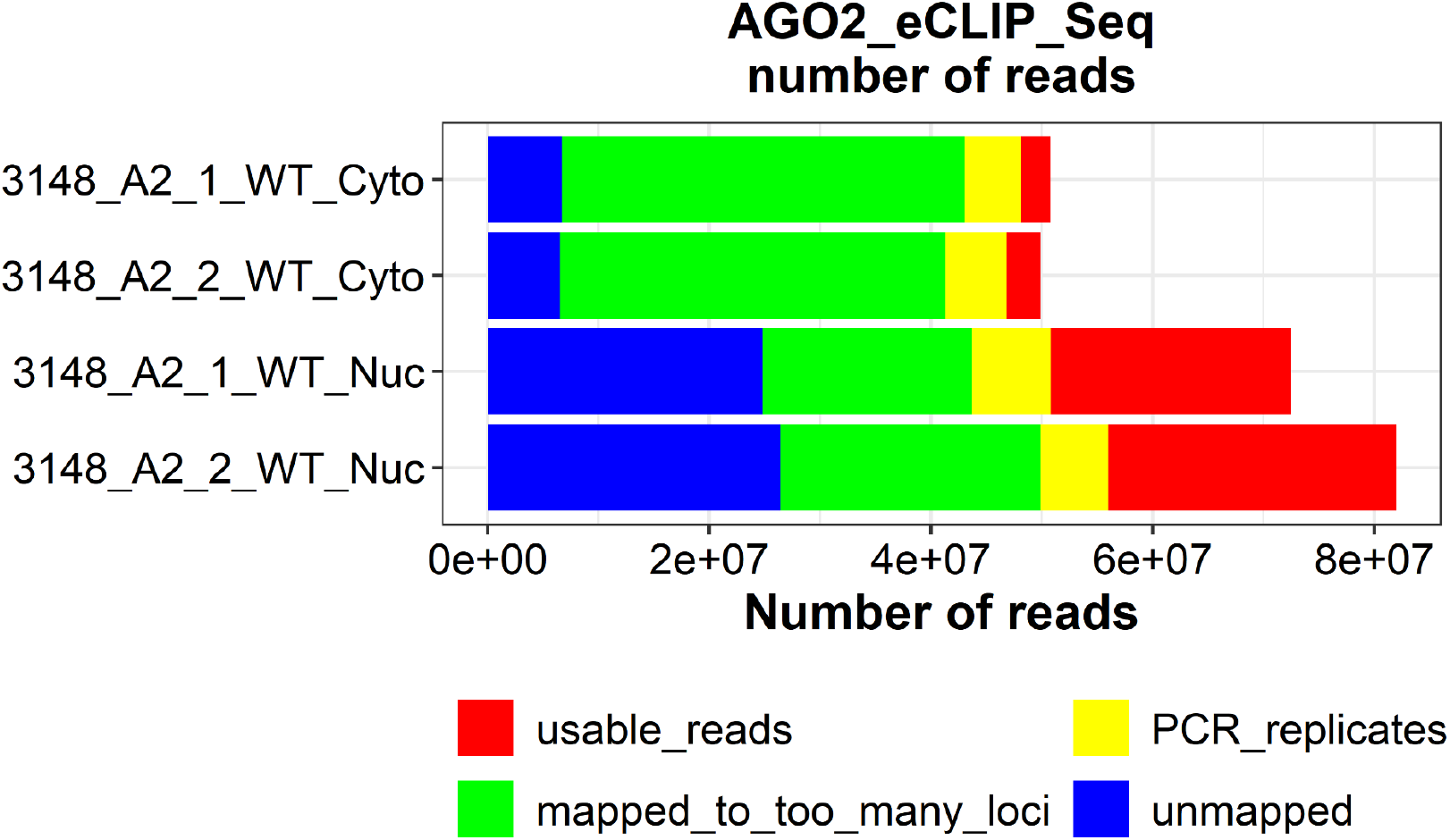
Anti-AGO2 eCLIP-seq nuclear samples. Sequencing reads distribution, showing the total number of usable reads.

**Supplemental Figure S2A.**
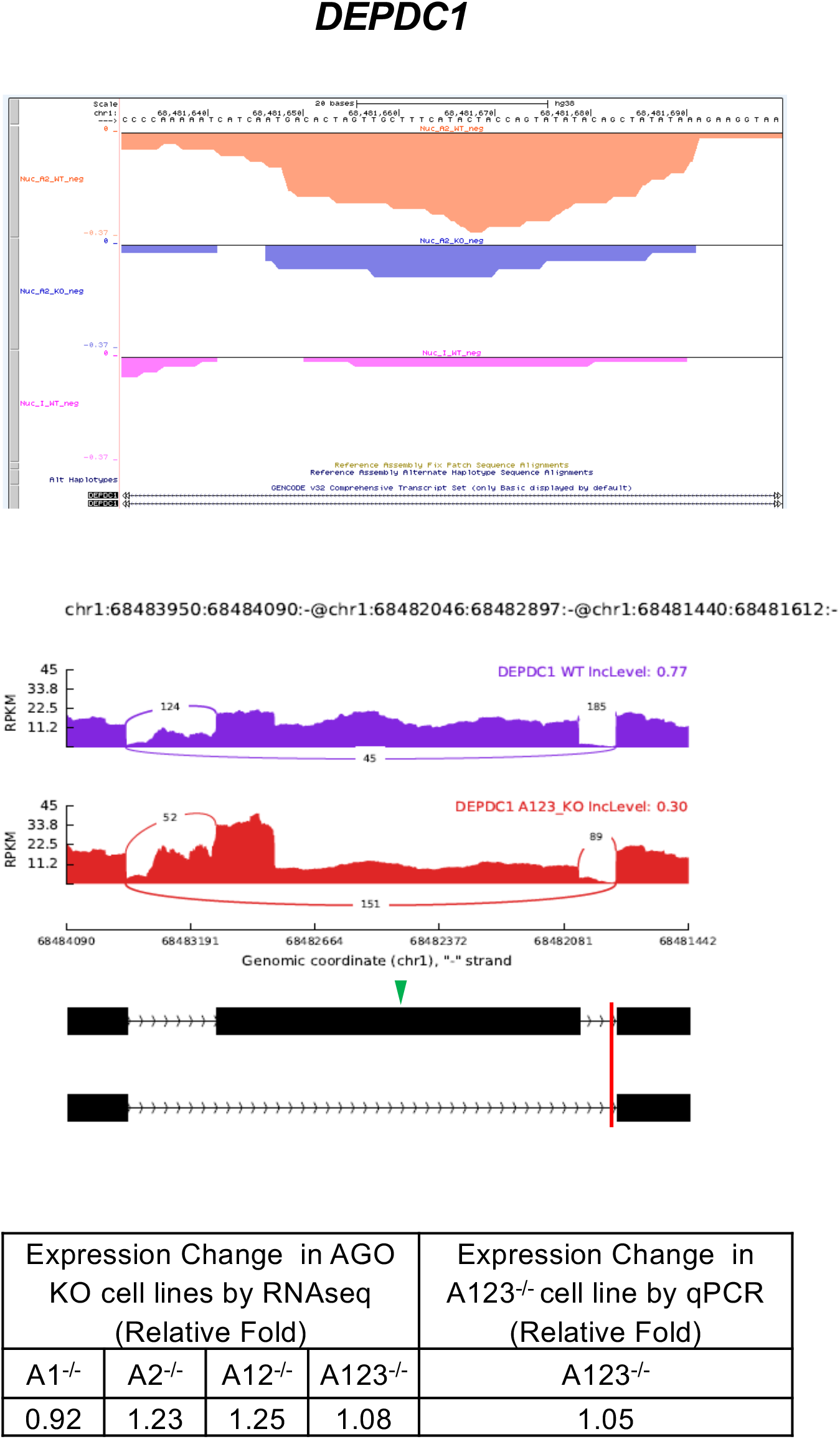
Splicing changes for a gene candidate in *AGO1/2/3* KO cells with AGO2 protein binding cluster: DEPDC1. *Top*. AGO2 protein binding clusters within *DEPDC1* identified by AGO2 eCLIP-seq. Orange: Wild type cells. Blue: *AGO2* knockout cells. Pink: Wild type input control. All clusters were located in skipped exon events nearby within intron. *Middle*. Sashimi plot for significant skipped exon event by RNA-seq analysis. Purple: Wild type cells. Red: *AGO1/2/3* knockout cells. Green arrowhead: excluded/included exon. Red vertical line: location of AGO2 binding cluster. For inclusion in analysis we required peaks to possess a *p* value <0.05 and a >4-fold enrichment in read number for wild-type verse AGO2 knockouts. *Bottom*. Expression change by RNA Seq and qPCR in AGO KO cell lines.

**Supplemental Figure S2B.**
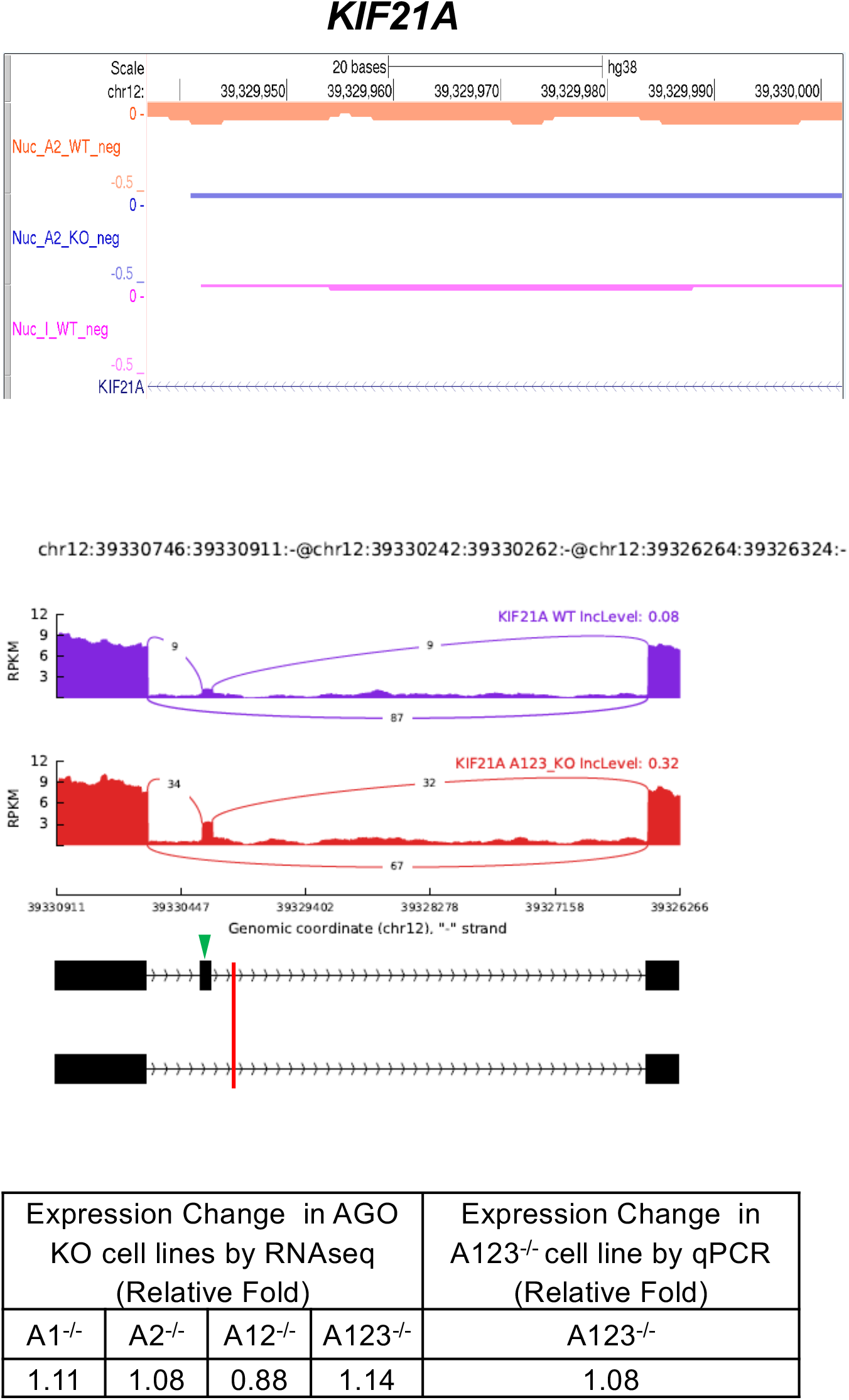
Splicing changes for a gene candidate in *AGO1/2/3 KO* cells with AGO2 protein binding cluster: KIF21A. *Top*. AGO2 protein binding clusters within *KIF21A* identified by AGO2 eCLIP-seq. Orange: Wild type cells. Blue: *AGO2* knockout cells. Pink: Wild type input control. All clusters were located in skipped exon events nearby within intron. *Middle*. Sashimi plot for significant skipped exon event by RNA-seq analysis. Purple: Wild type cells. Red: *AGO1/2/3* knockout cells. Green arrowhead: excluded/included exon. Red vertical line: location of AGO2 binding cluster. For inclusion in analysis we required peaks to possess a *p* value <0.05 and a >4-fold enrichment in read number for wild-type verse AGO2 knockouts. *Bottom*. Expression change by RNA Seq and qPCR in AGO KO cell lines.

**Supplemental Figure S2C.**
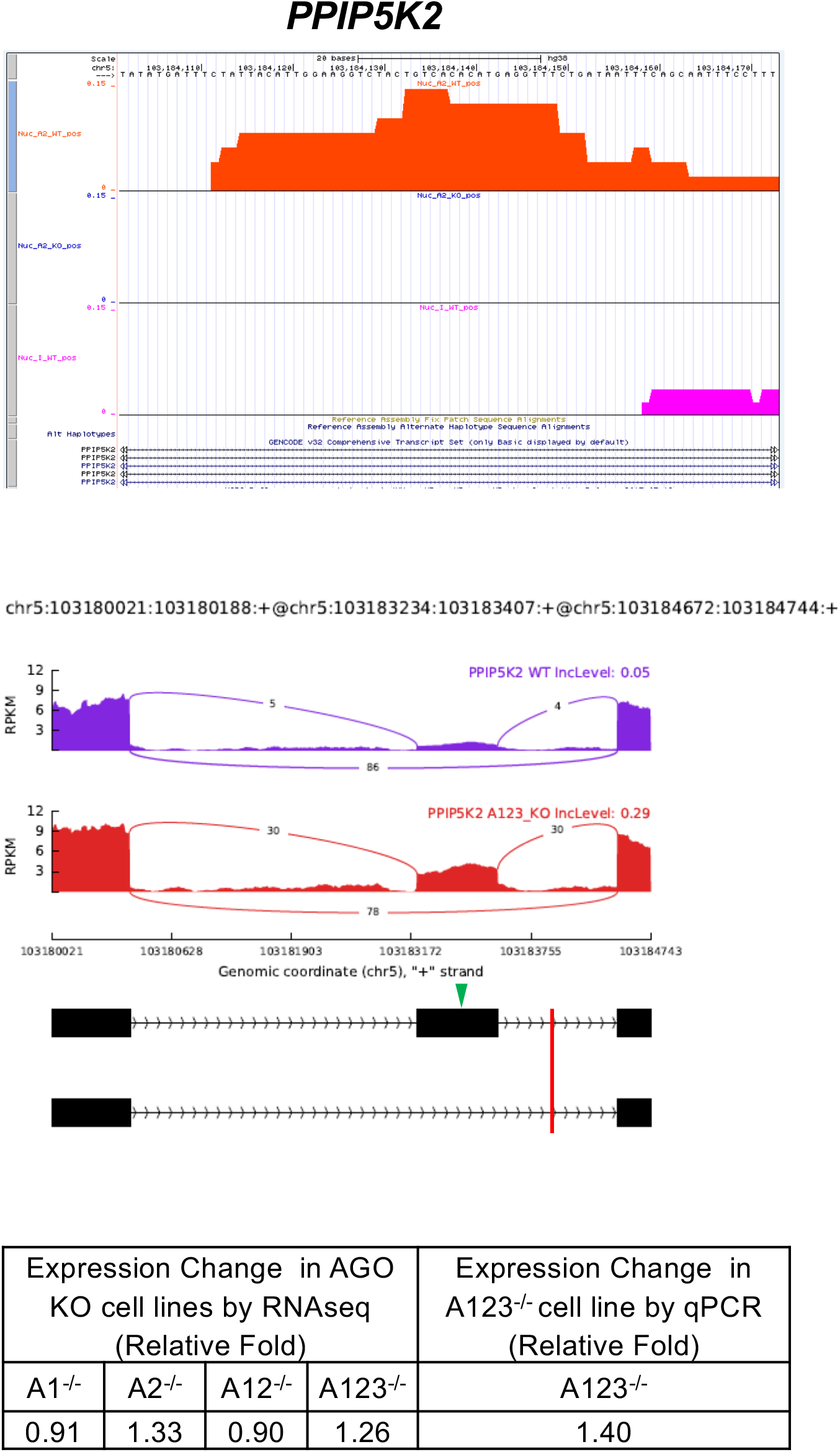
Splicing changes for a gene candidate in *AGO1/2/3* KO cells with AGO2 protein binding cluster: PPIP5K2. *Top*. AGO2 protein binding clusters within *PPIP5K2* identified by AGO2 eCLIP-seq. Red: Wild type cells. Navy: *AGO2* knockout cells. Pink: Wild type input control. All clusters were located in skipped exon events nearby within intron. *Middle*. Sashimi plot for significant skipped exon event by RNA-seq analysis. Purple: Wild type cells. Red: *AGO1/2/3* knockout cells. Green arrowhead: excluded/included exon. Red vertical line: location of AGO2 binding cluster. For inclusion in analysis we required peaks to possess a *p* value <0.05 and a >4-fold enrichment in read number for wild-type verse *AGO2* knockouts. *Top*. Expression change by RNA Seq and qPCR in AGO KO cell lines.

**Supplemental Figure SD.**
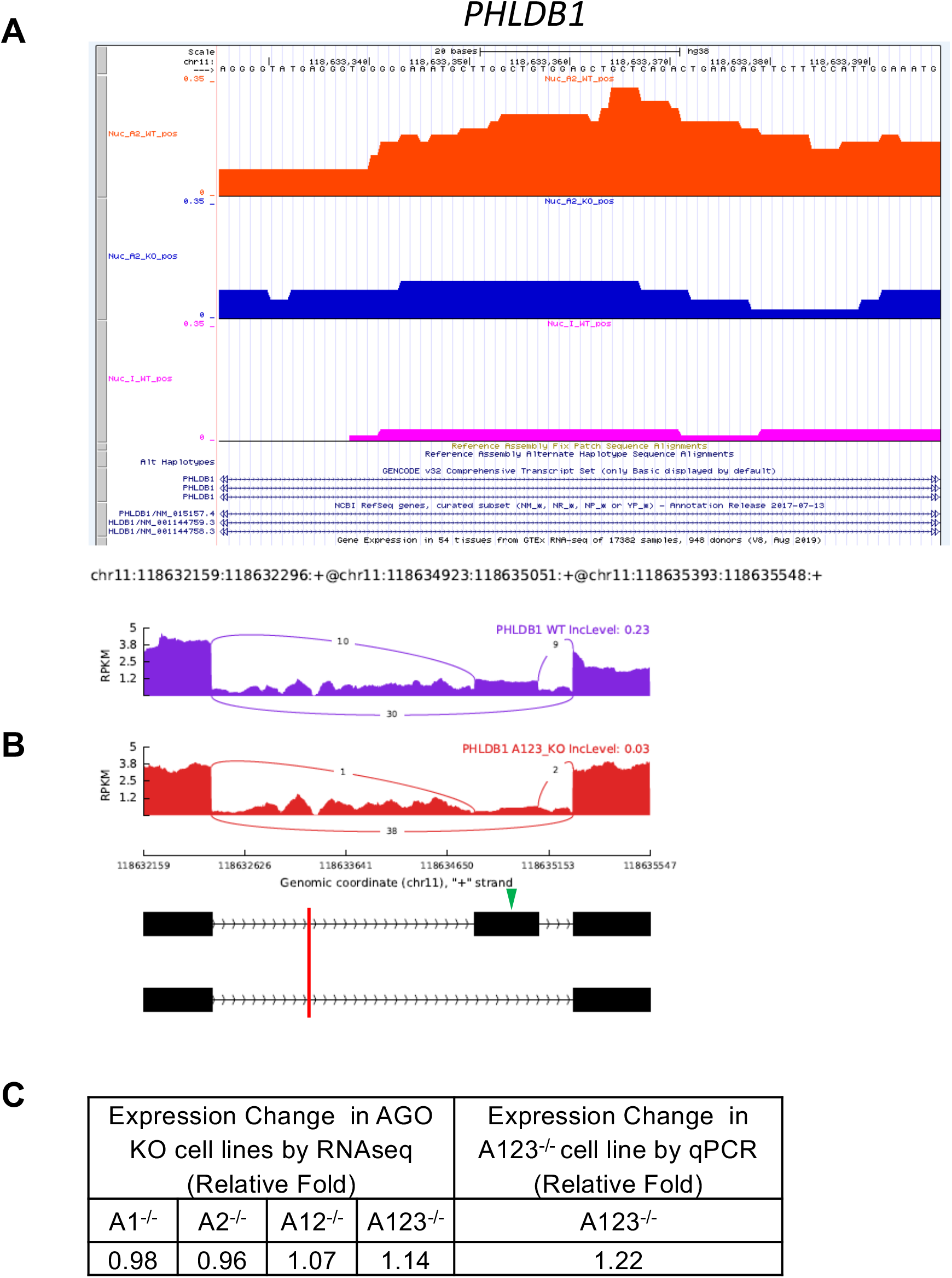
Splicing changes for a gene candidate in *AGO1/2/3* KO cells with AGO2 protein binding cluster: PHLDB1. *Top*. AGO2 protein binding clusters within *PHLDB1* identified by AGO2 eCLIP-seq. Red: Wild type cells. Navy: *AGO2* knockout cells. Pink: Wild type input control. All clusters were located in skipped exon events nearby within intron. *Middle*. Sashimi plot for significant skipped exon event by RNA-seq analysis. Purple: Wild type cells. Red: *AGO1/2/3* knockout cells. Green arrowhead: excluded/included exon. Red vertical line: location of AGO2 binding cluster. For inclusion in analysis we required peaks to possess a *p* value <0.05 and a >4-fold enrichment in read number for wild-type verse AGO2 knockouts. *Bottom*. Expression change by RNA Seq and qPCR in AGO KO cell lines.

**Supplemental Figure S2E.**
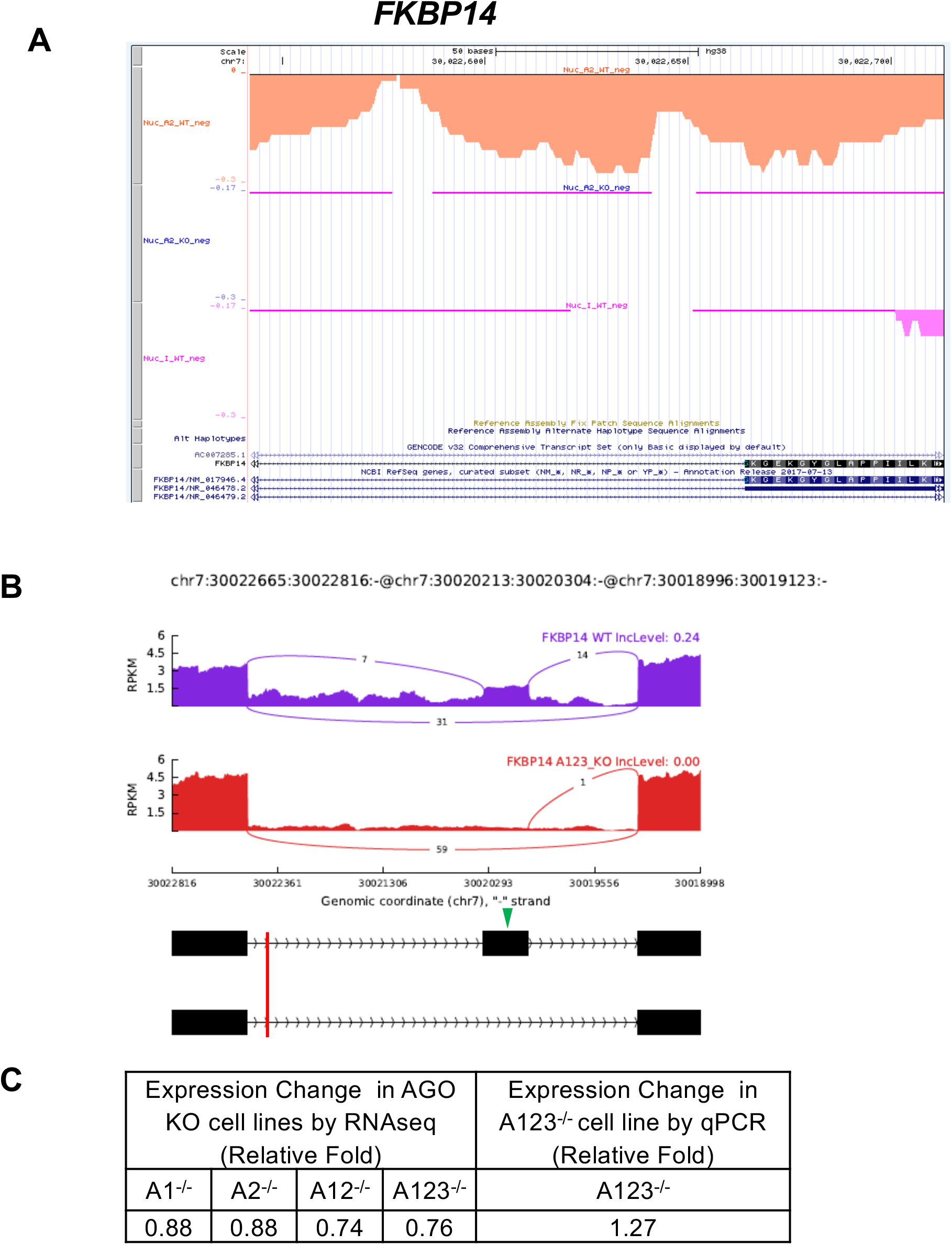
Splicing changed gene candidate in *AGO1/2/3* KO cells with AGO2 binding cluster: *FKBP14*. *Top*. AGO2 protein binding clusters within *FKBP14* identified by AGO2 eCLIP-seq. Orange: Wild type cells. Blue: *AGO2* knockout cells. Pink: Wild type input control. All clusters were located in skipped exon events nearby within intron. *Middle*. Sashimi plot for significant skipped exon event by RNA-seq analysis. Purple: Wild type cells. Red: *AGO1/2/3* knockout cells. Green arrowhead: excluded/included exon. Red vertical line: location of AGO2 binding cluster. For inclusion in analysis we required peaks to possess a *p* value <0.05 and a >4-fold enrichment in read number for wild-type verse *AGO2* knockouts. *Bottom*. Expression change by RNA Seq and qPCR in AGO KO cell lines.

**Supplemental Figure S2F.**
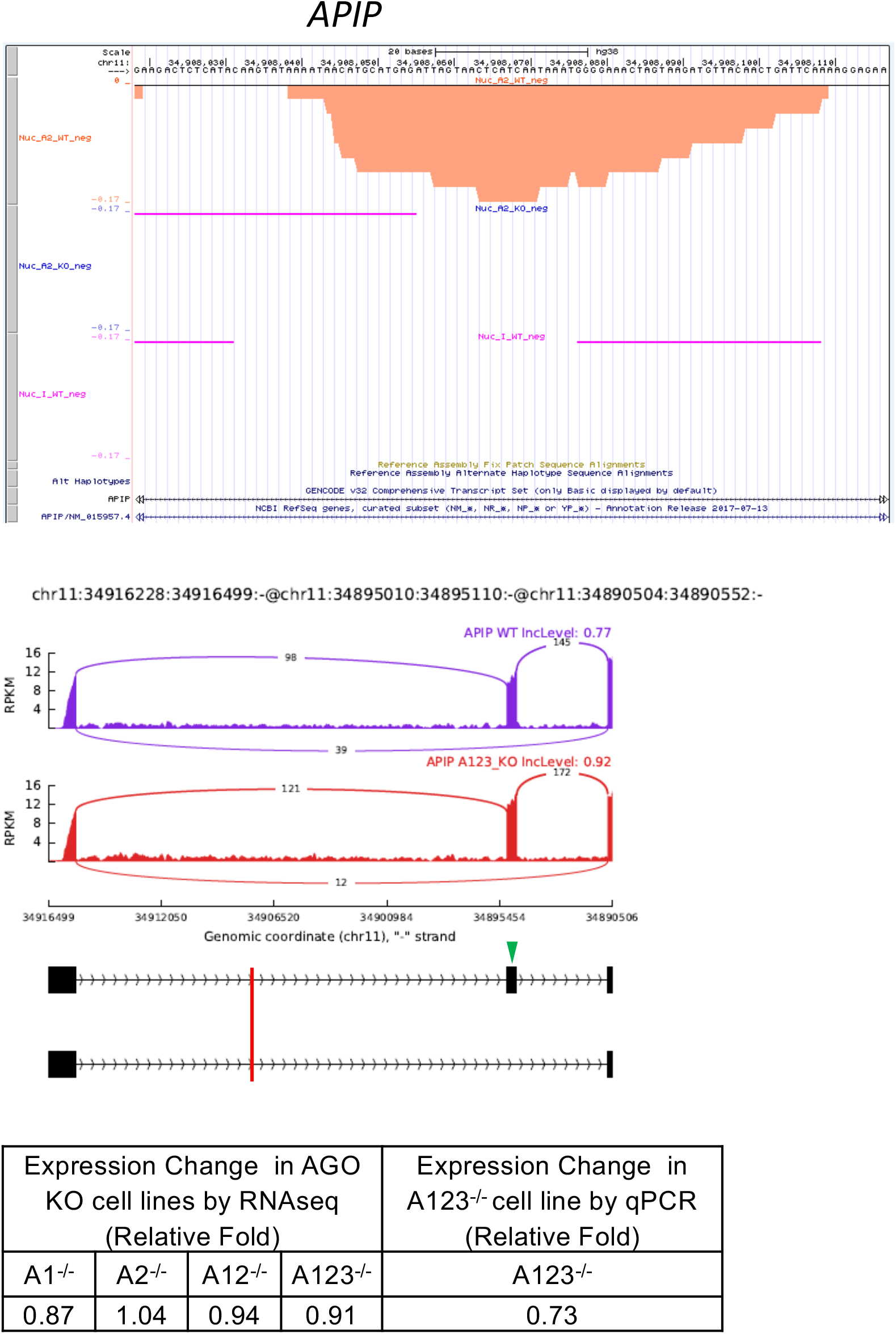
Splicing changed gene candidate in *AGO1/2/3* KO cells with AGO2 protein binding cluster: *APIP*. *Top*. AGO2 protein binding clusters within *APIP* identified by AGO2 eCLIP-seq. Orange: Wild type cells. Blue: *AGO2* knockout cells. Pink: Wild type input control. All clusters were located in skipped exon events nearby within intron. *Middle*. Sashimi plot for significant skipped exon event by RNA-seq analysis. Purple: Wild type cells. Red: *AGO1/2/3* knockout cells. Green arrowhead: excluded/included exon. Red vertical line: location of AGO2 binding cluster. For inclusion in analysis we required peaks to possess a *p* value <0.05 and a >4-fold enrichment in read number for wild-type verse *AGO2* knockouts. *Bottom*. Expression change by RNA Seq and qPCR in AGO KO cell lines.

**Supplemental Figure S2G.**
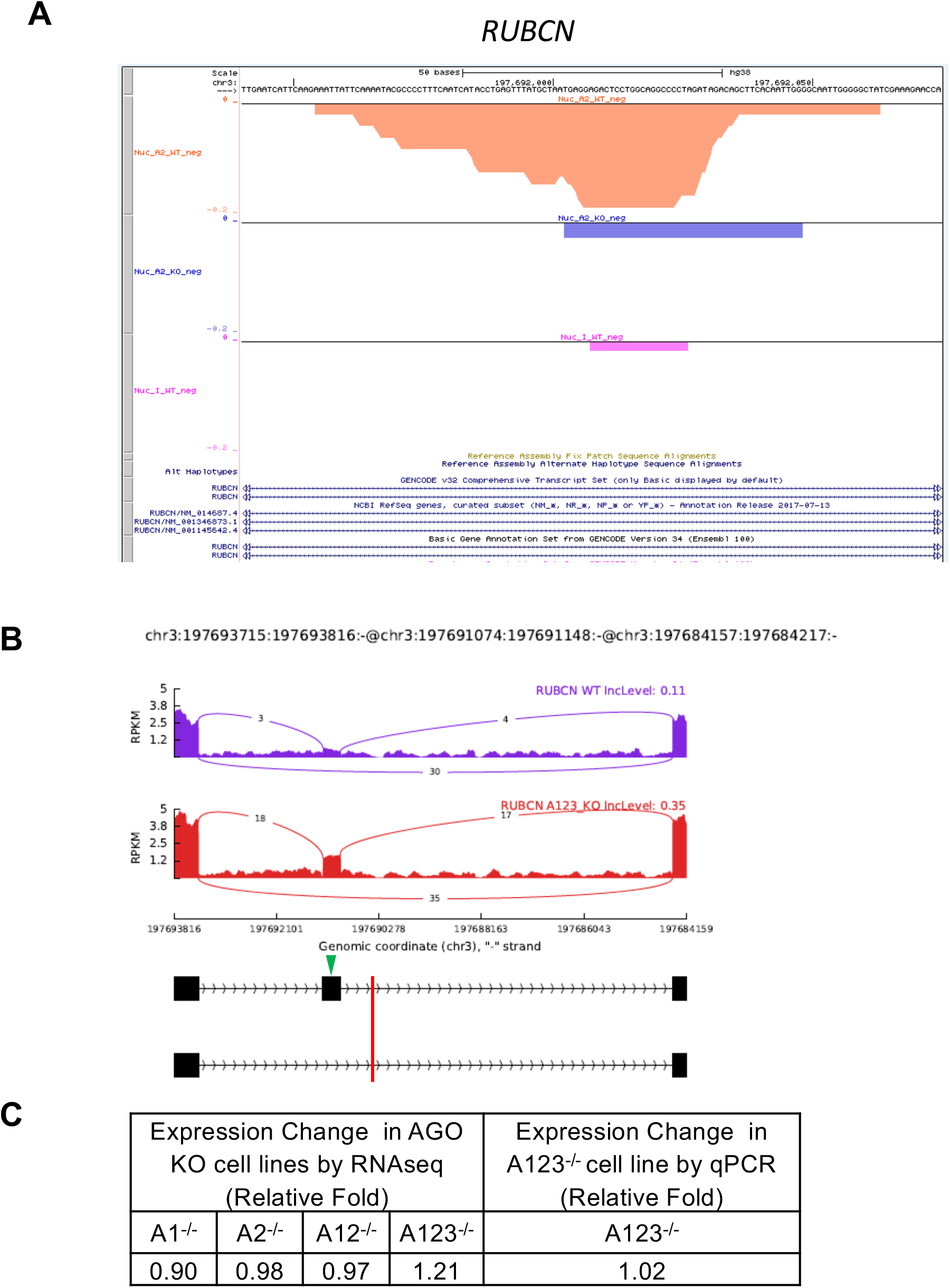
Splicing changed gene candidate in *AGO1/2/3* KO cells with AGO2 protein binding cluster: *RUBCN*. *Top*. AGO2 protein binding clusters within *RUBCN* identified by AGO2 eCLIP-seq. Orange: Wild type cells. Blue: *AGO2* knockout cells. Pink: Wild type input control. All clusters were located in skipped exon events nearby within intron. *Middle*. Sashimi plot for significant skipped exon event by RNA-seq analysis. Purple: Wild type cells. Red: *AGO1/2/3* knockout cells. Green arrowhead: excluded/included exon. Red vertical line: location of AGO2 binding cluster. For inclusion in analysis we required peaks to possess a *p* value <0.05 and a >4-fold enrichment in read number for wild-type verse *AGO2* knockouts. *Bottom*. Expression change by RNA Seq and qPCR in AGO KO cell lines.

**Supplemental Figure S2H.**
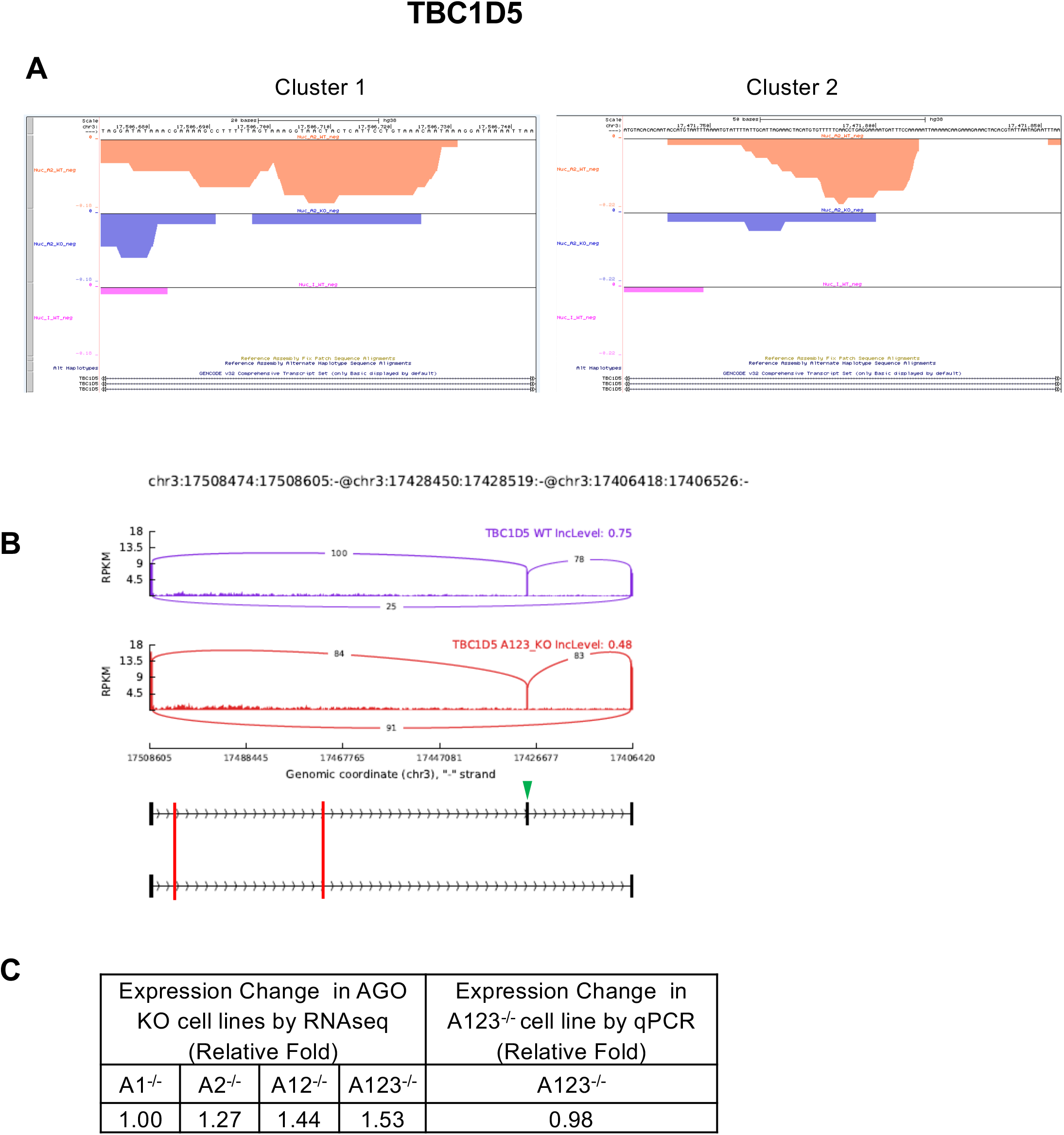
Splicing changed gene candidate in *AGO1/2/3* KO cells with AGO2 protein binding cluster: *TBC1D5*. *Top*. AGO2 protein binding clusters within *TBC1D5* identified by AGO2 eCLIP-seq. Orange: Wild type cells. Blue: *AGO2* knockout cells. Pink: Wild type input control. All clusters were located in skipped exon events nearby within intron. *Middle*. Sashimi plot for significant skipped exon event by RNA-seq analysis. Purple: Wild type cells. Red: *AGO1/2/3* knockout cells. Green arrowhead: excluded/included exon. Red vertical line: location of AGO2 binding cluster. For inclusion in analysis we required peaks to possess a *p* value <0.05 and a >4-fold enrichment in read number for wild-type verse *AGO2* knockouts. *Bottom*. Expression change by RNA Seq and qPCR in AGO KO cell lines.

**Supplemental Figure S3, related to Figure 5.**
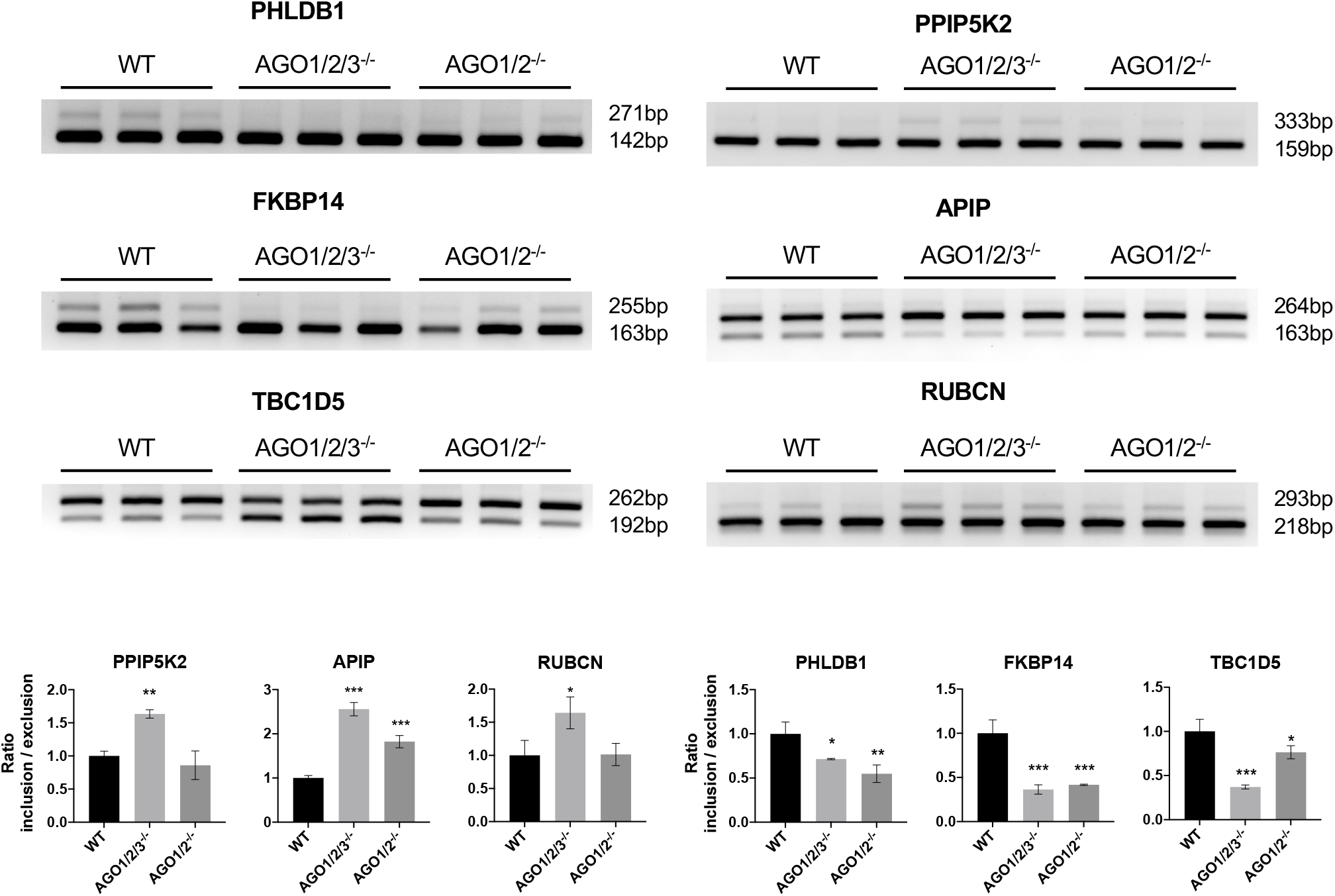
Semiquantitative PCR validation of skipped exon events in *AGO1/2* KO and *AGO1/2/3* KO cells by different PCR primer sets. Semiquantitative PCR validation of skipped exon events in *AGO1/2/3* KO and *AGO1/2* KO cells. Error bars represent standard deviation (SD). *P < 0.05; **P < 0.01; ***P < 0.001 compared with WT by one-way ANOVA and Dunnett’s multiple comparisons test.

**Supplemental Figure S4.**
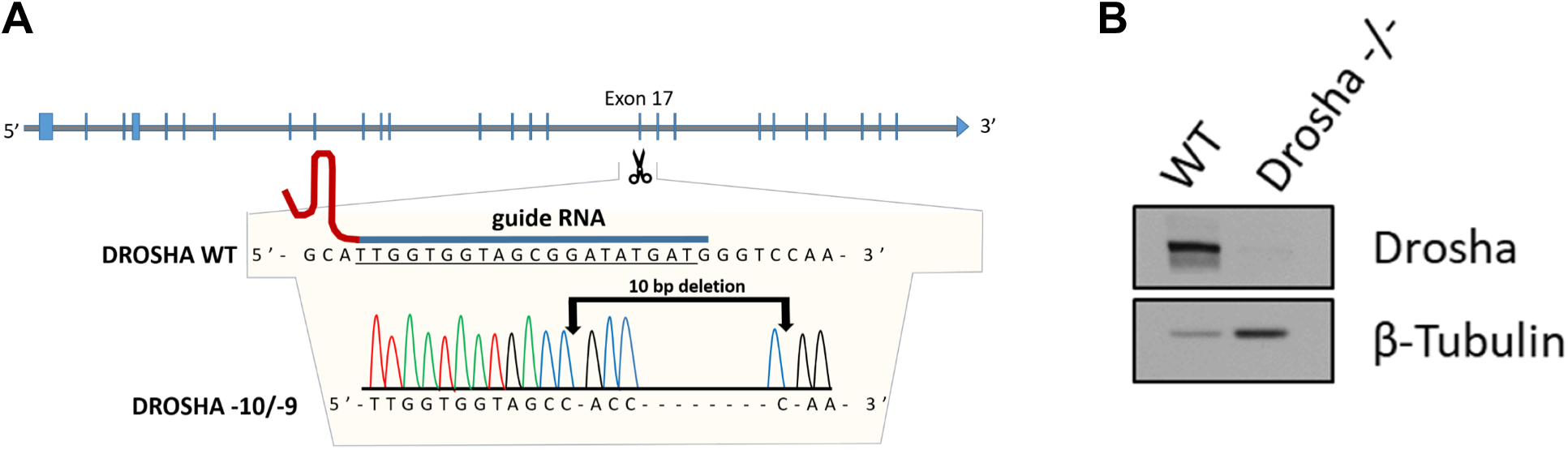
*DROSHA* KO cell. (*A*) Location of guide RNA and deletions from CRISPR/Cas9-derived HCT116 *DROSHA* knockout cell line. (*B*) Western blot validating the *DROSHA* −10/−9 cell line.

**Supplemental Figure S5, related to Figure 6.**
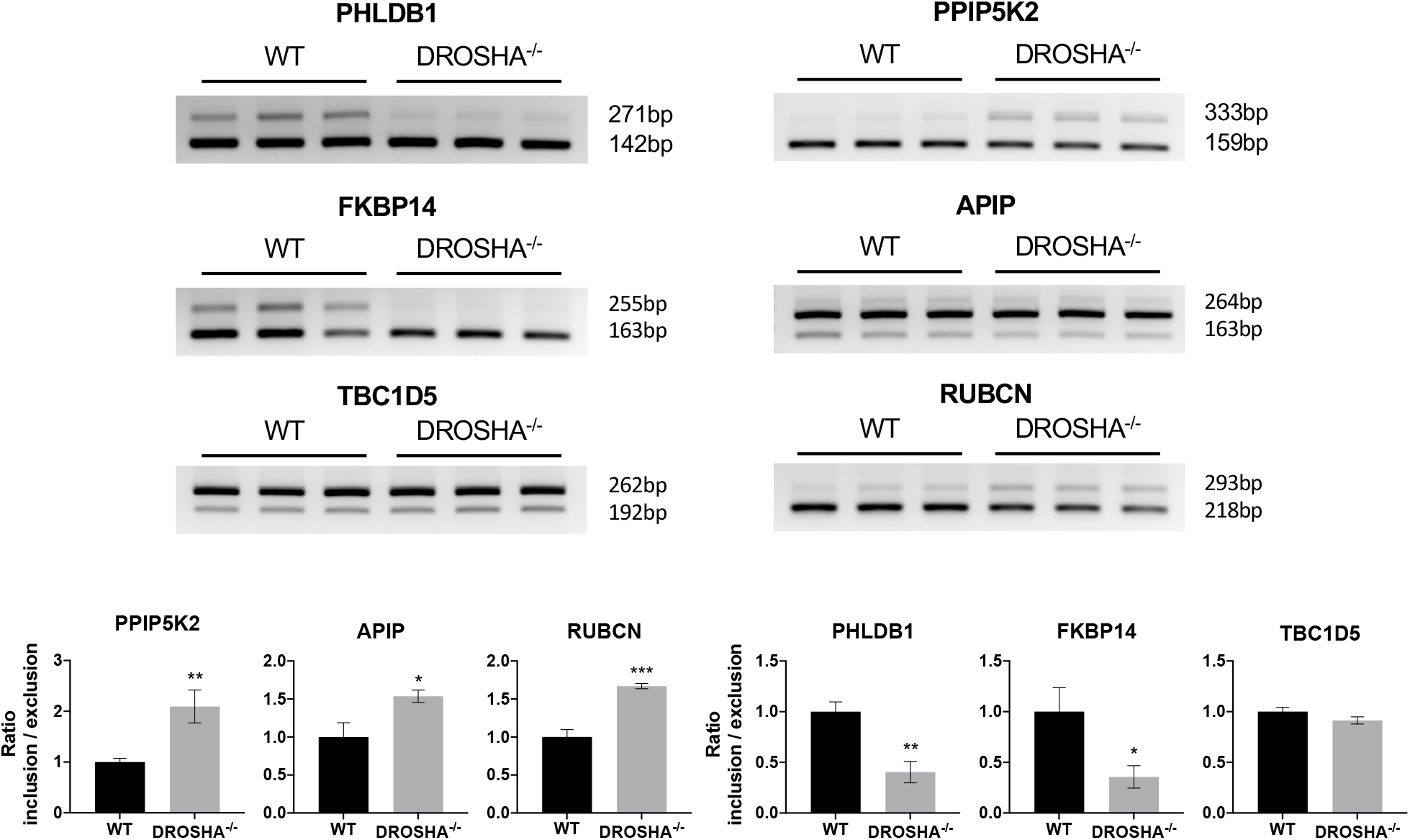
Semiquantitative PCR validation of skipped exon events in Drosha KO cells by different PCR primer sets. Skipped exon events in *DROSHA* KO cells. Semiquantitative PCR validation of skipped exon events in *DROSHA* KO cells. Error bars represent standard deviation (SD). *P < 0.05; **P < 0.01; ***P < 0.001 compared with WT by t-test.

**Supplemental Figure S6, related to Figure 7.**
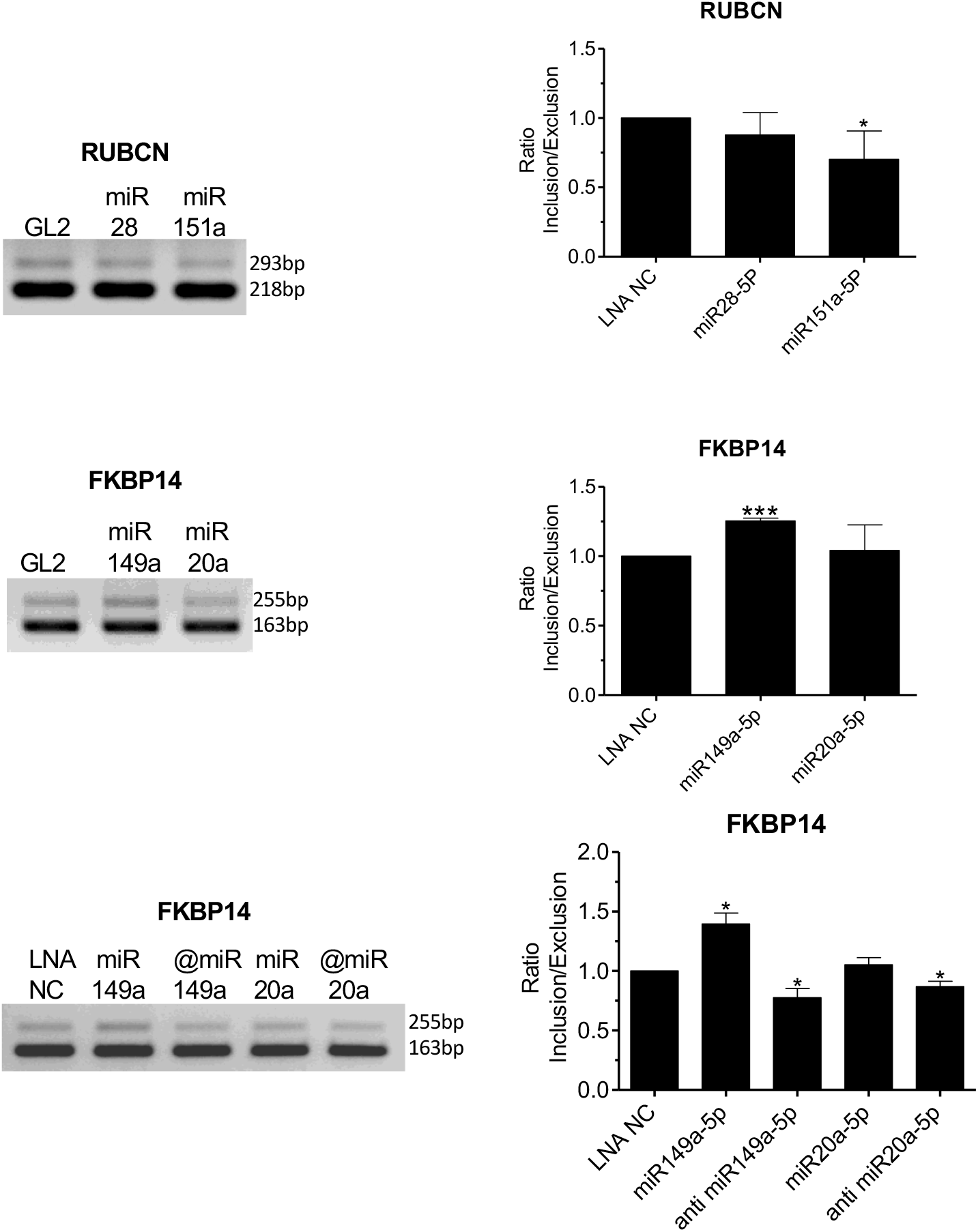
Semiquantitative agarose electrophoresis validation of skipped exon events after treated by miRNA mimics (50 nM) and LNA anti-miR oligonucleotides (50 nM). Skipped exon events in WT cells. Error bars represent standard deviation (SD). *P < 0.05; **P < 0.01; ***P < 0.001 compared with WT by t-test.

**Supplementary Table S1A.**
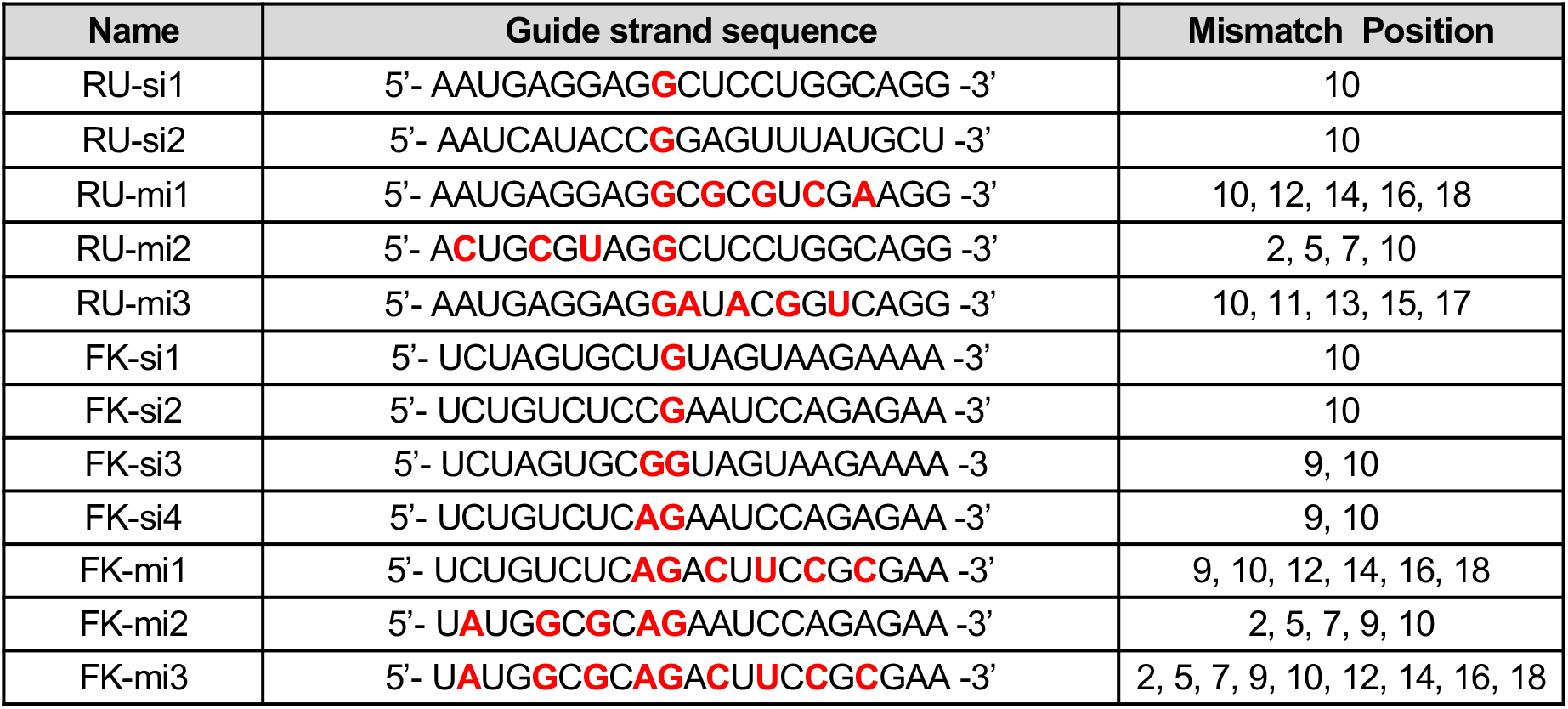
Double strand RNA sequences and mismatch position.

**Supplementary Table S1B.**
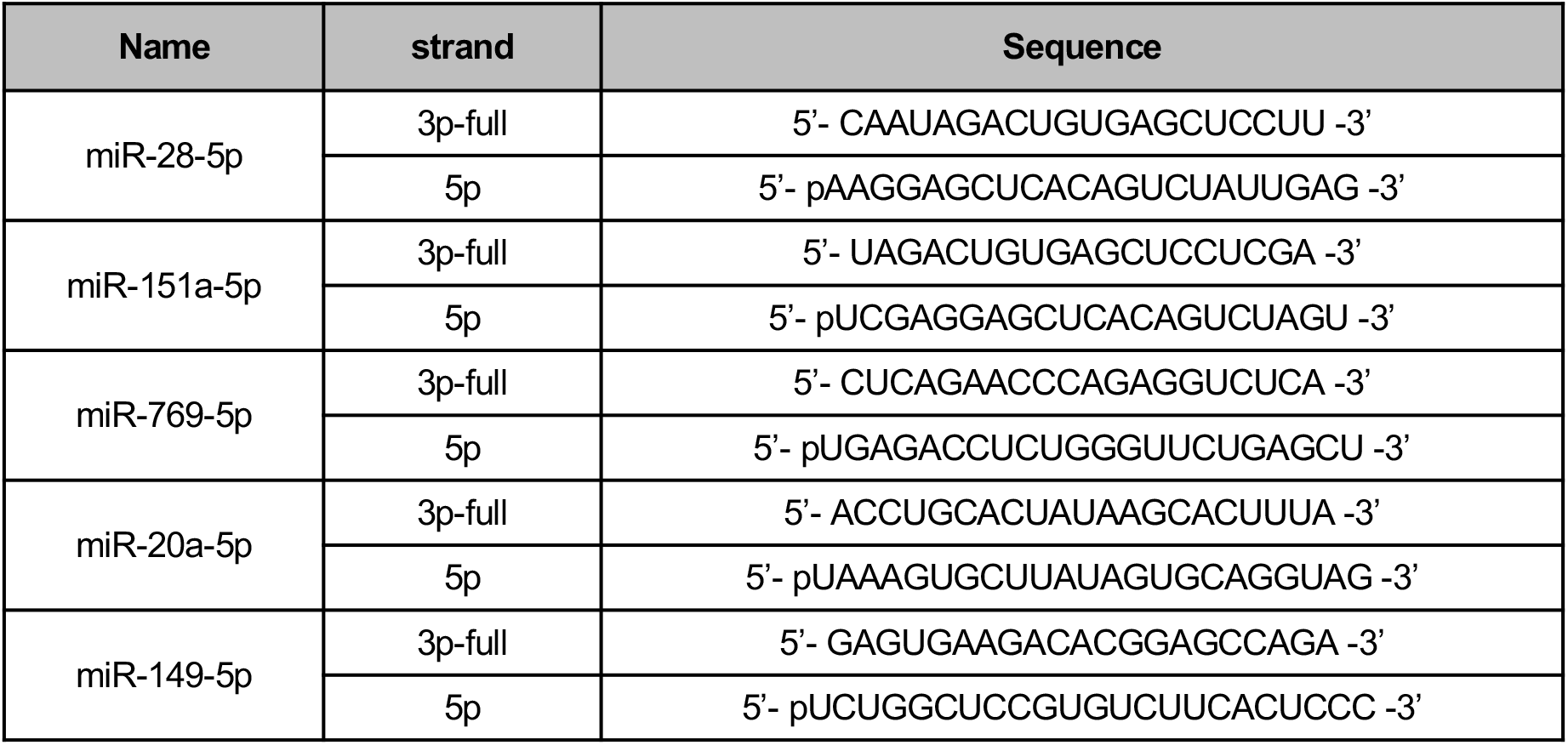
miRNA mimic sequence.

**Supplementary Table S2.**
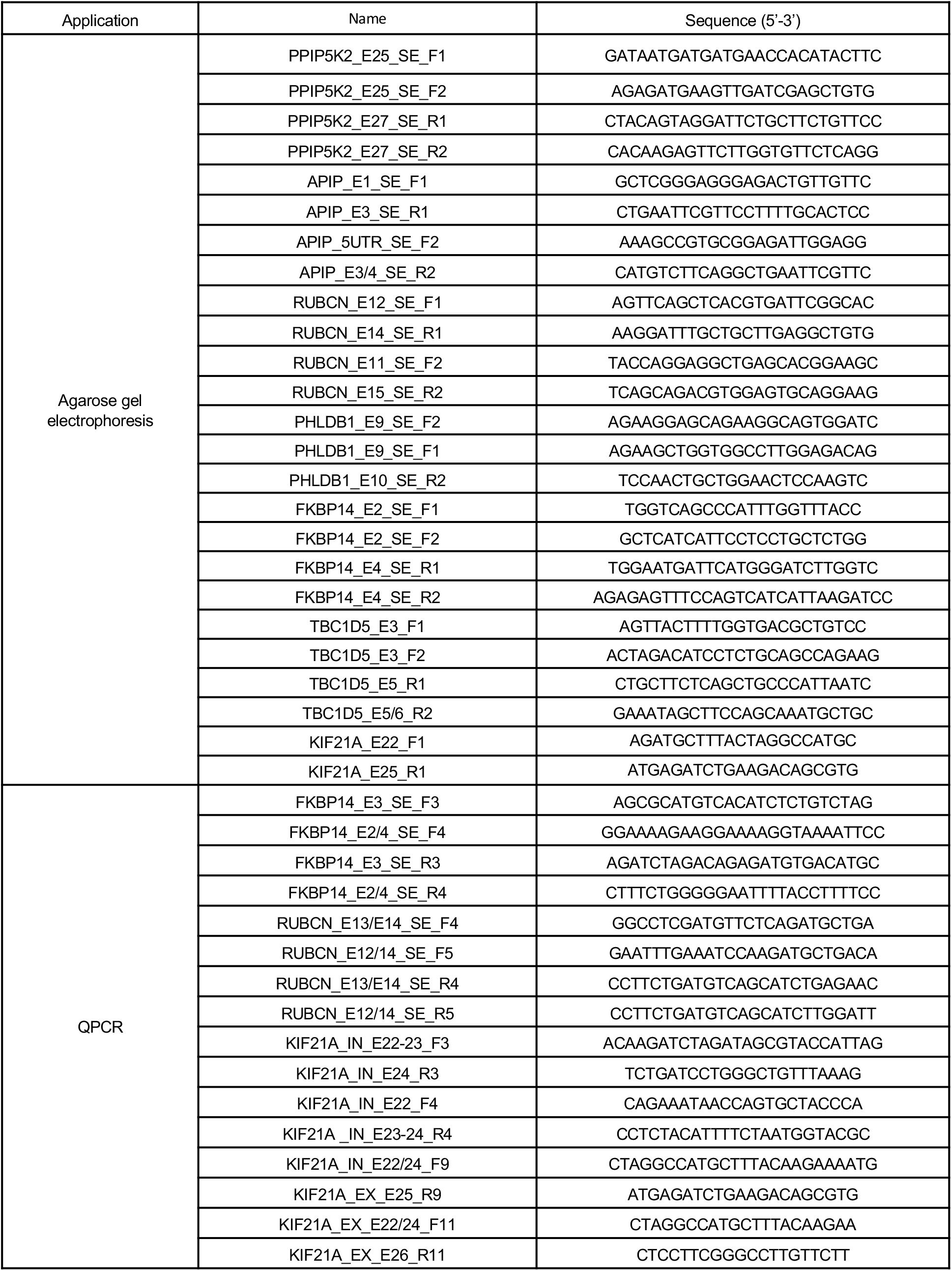
PCR primers for gel electrophoresis and qPCR analysis.

**Supplementary Table S3.**
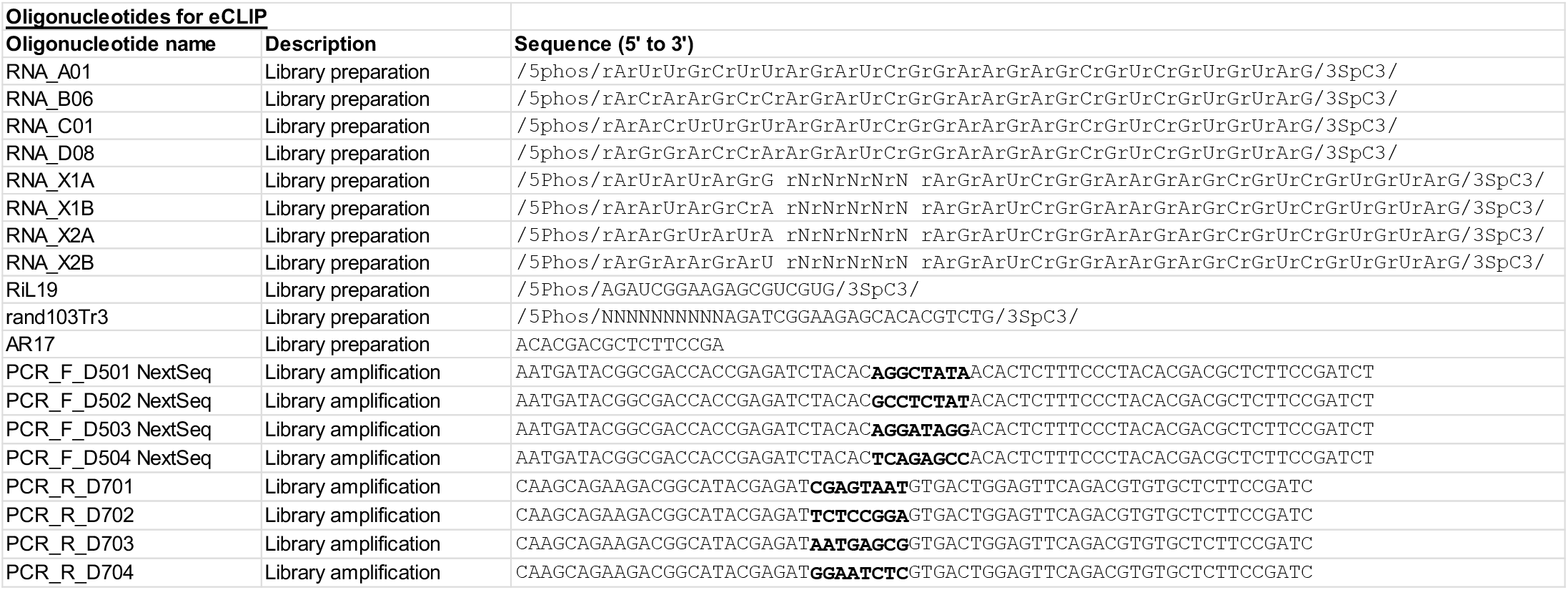
Oligonucleotides used in eCLIP library preparation

